# A fine-tuned vector-parasite dialogue in tsetse’s cardia determines peritrophic matrix integrity and trypanosome transmission success

**DOI:** 10.1101/207993

**Authors:** Aurélien Vigneron, Emre Aksoy, Brian L Weiss, Xiaoli Bing, Xin Zhao, Erick O Awuoche, Michelle O’Neill, Yineng Wu, Geoffrey M Attardo, Serap Aksoy

## Abstract

Arthropod vectors have multiple physical and immunological barriers that impede the development and transmission of parasites to new vertebrate hosts. These barriers include the peritrophic matrix (PM), a chitinous barrier that separates the blood bolus from the midgut epithelia and inturn, modulates vector-microbiota interactions. In tsetse flies, a sleeve-like PM is continuously produced by the cardia organ located at the fore- and midgut junction. African trypanosomes, *Trypanosoma brucei*, must bypass the PM twice; first to colonize the midgut and secondly to reach the salivary glands (SG), to complete their transmission cycle in tsetse. However, not all flies with midgut infections develop mammalian transmissible SG infections - the reasons for which are unclear. Here, we used transcriptomics, microscopy and functional genomics analyses to understand the factors that regulate parasite migration from midgut to SG. In flies with midgut infections only, parasites fail to cross the PM as they are eliminated from the cardia by reactive oxygen intermediates (ROIs) - albeit at the expense of collateral cytotoxic damage to the cardia. In flies with midgut and SG infections, expression of genes encoding components of the PM is reduced in the cardia, and structural integrity of the PM barrier is compromised. Under these circumstances trypanosomes traverse through the newly secreted and compromised PM. The process of PM attrition that enables the parasites to re-enter into the midgut lumen is apparently mediated by components of the parasites residing in the cardia. Thus, a fine-tuned dialogue between tsetse and trypanosomes at the cardia determines the outcome of PM integrity and trypanosome transmission success.

**Author summary:** Insects are responsible for transmission of parasites that cause deadly diseases in humans and animals. Understanding the key factors that enhance or interfere with parasite transmission processes can result in new control strategies. Here, we report that a proportion of tsetse flies with African trypanosome infections in their midgut can prevent parasites from migrating to the salivary glands, albeit at the expense of collateral damage. In a subset of flies with gut infections, the parasites manipulate the integrity of a midgut barrier, called the peritrophic matrix, and reach the salivary glands for transmission to the next mammal. Either targeting parasite manipulative processes or enhancing peritrophic matrix integrity could reduce parasite transmission.

## Introduction

Insects are essential vectors for the transmission of microbes that cause devastating diseases in humans and livestock. Many of these diseases lack effective vaccines and drugs for control in mammalian hosts. Hence, reduction of insect populations, as well as approaches that reduce the transmission efficiency of pathogens by insect vectors, are explored for disease control. Tsetse flies transmit African trypanosomes, which are the causative agents of human and animal African trypanosomiases. These diseases can be fatal if left untreated and inflict significant socio economic hardship across a wide swath of sub-Saharan Africa [1, 2]. The phenomenon of antigenic variation the parasite displays in its mammalian host has prevented the development of vaccines, and easily administered and affordable drugs are unavailable. However, tsetse population reduction can significantly curb disease, especially during times of endemicity [3, 4]. In addition, strategies that reduce parasite transmission efficiency by the tsetse vector can prevent disease emergence. A more complete understanding of parasite-vector dynamics is essential for the development of such control methods.

For transmission to new vertebrate hosts, vector-borne parasites have to first successfully colonize their respective vectors. This requires that parasites circumvent several physical and immune barriers as they progress through their development in the vector. One prominent barrier they face in the midgut is the peritrophic matrix (PM), which is a chitinous, proteinaceous structure that separates the epithelia from the blood meal [5–7]. In Anopheline mosquitoes the presence of the PM benefits the vector by regulating the commensal gut microbiota and preventing pathogens from invading the hemocoel [8]. In tsetse and sand flies, the PM plays a crucial role as an infection barrier by blocking parasite development and colonization [9, 10]. The presence of the PM can also be exploited by microbes to promote their survival in the gut lumen. The agent of Lyme disease, *Borrelia burgdorferi*, binds to the tick vector’s gut and exploits the PM for protection from the harmful effects of blood-filled gut lumen [11]. Unlike vectors that produce a type I PM in response to blood feeding, tsetse’s sleeve-like type II PM is constitutively produced by the cardia organ, which is located at the junction of the fore- and midgut. Upon entering the gut lumen, long-slender bloodstream form (BSF) trypanosomes (*Trypanosoma brucei*) are lysed while short-slender BSFs differentiate to midgut-adapted procyclic forms (PCF) [12]. During these lysis and differentiation processes, BSF parasites shed their dense coat composed of Variant Surface Glycoproteins (VSGs) into the midgut environment [12]. These molecules are then internalized by cells in the cardia, where they transiently inhibit the production of a structurally robust PM. This process promotes infection establishment by enabling trypanosomes to traverse the PM barrier and invade the midgut ectoperitrophic space (ES) [9]. After entering the ES, trypanosomes face strong epithelial immune responses, which hinder parasite gut colonization success. Detection of PCF parasites in the ES induces the production of trypanocidal antimicrobial peptides [13, 14], reactive oxygen intermediates (ROIs) [15], PGRP-LB [16] and tsetse-EP protein [17]. A combination of these immune effectors eliminate trypanosomes from the majority of flies, leaving only a small percentage of flies infected with PCF parasites in their midgut. The PCF parasites move to tsetse’s cardia where they differentiate into long and short epimastigote forms. These cells then cross the PM for the second time to enter back into the fly’s gut lumen and migrate through the foregut into the salivary glands (SG) for further differentiation into mammalian infective metacyclic forms [18, 19]. Interestingly, the SG infection process, which is necessary for disease transmission, succeeds in only a subset of flies with midgut infections [20]. Even though midgut trypanosomes fail to colonize tsetse’s SG in a subset of flies, parasites persist in the midgut for the remainder of the fly’s adult life. The physiological barriers that prevent SG colonization in the subset of midgut only infected flies remain unknown.

In this study, we investigated the molecular and cellular mechanisms that prevent parasites from colonizing the SG in a subset of flies with successful midgut infections. Our results show a robust host oxidative stress response reduces parasite survival in the cardia. While preventing parasites from further development, this immune response is costly for tsetse’s cellular integrity and results in extensive damage to cardia tissues. In contrast, less cellular damage is observed in the cardia of flies with midgut parasites that give rise to SG infections. Our results indicate that the ability of the parasites to successfully bypass the PM barrier in the cardia is essential for the establishment of SG infections. We discuss the molecular interactions that regulate this complex and dynamic vector-parasite relationship in the cardia organ, an essential regulator of disease transmission.

## Results and Discussion

### Trypanosome infection dynamics in tsetse

Tsetse display strong resistance to infection with trypanosomes. By 3-6 days post acquisition (dpa), parasites that have entered into the ES of the midgut are eliminated by induced vector immune responses from the majority of flies. When newly emerged *Glossina morsitans* adults (termed teneral) are provided with an infectious bloodmeal in their first feeding, midgut infection success is typically around 30-40% [21, 22]. However, in mature adults that have had at least one prior normal bloodmeal, the infection rate is lower, with only 1-5% of flies housing midgut infections [23, 24]. PCF parasites in susceptible flies replicate in the ES and move forward to the cardia where they differentiate into long and short epimastigote forms. About 6-10 dpa, parasites in the cardia re-enter the gut lumen and migrate through the foregut into the SG, where they differentiate to mammalian infective metacyclic forms within 20-30 days [18, 19].

To generate a suitable sample size of infected flies for down-stream experiments, we provided three groups of teneral *G. m. morsitans* adult females independent blood meals containing BSF trypanosomes (*Trypanosoma brucei brucei* RUMP 503) and obtained midgut infection rates of about 30% when microscopically analyzed 40 dpa (Fig 1A). When we analyzed the SG infection status of these gut infected flies, we detected SG infections in about 65% of individuals (Fig 1B). We chose the 40-day time point to accurately score midgut and SG infections, as in our experimental system and insectary environment SG infection status becomes accurately verifiable by microscopy at 30 dpa at the earliest. Hence, two forms of fly infections exist: non-permissive infections where parasites are restricted exclusively to the gut (hereafter designated ‘inf+/-’), and permissive infections where parasites are present in the gut and SGs (hereafter designated ‘inf+/+’) (Fig 1C).

**Fig 1.**
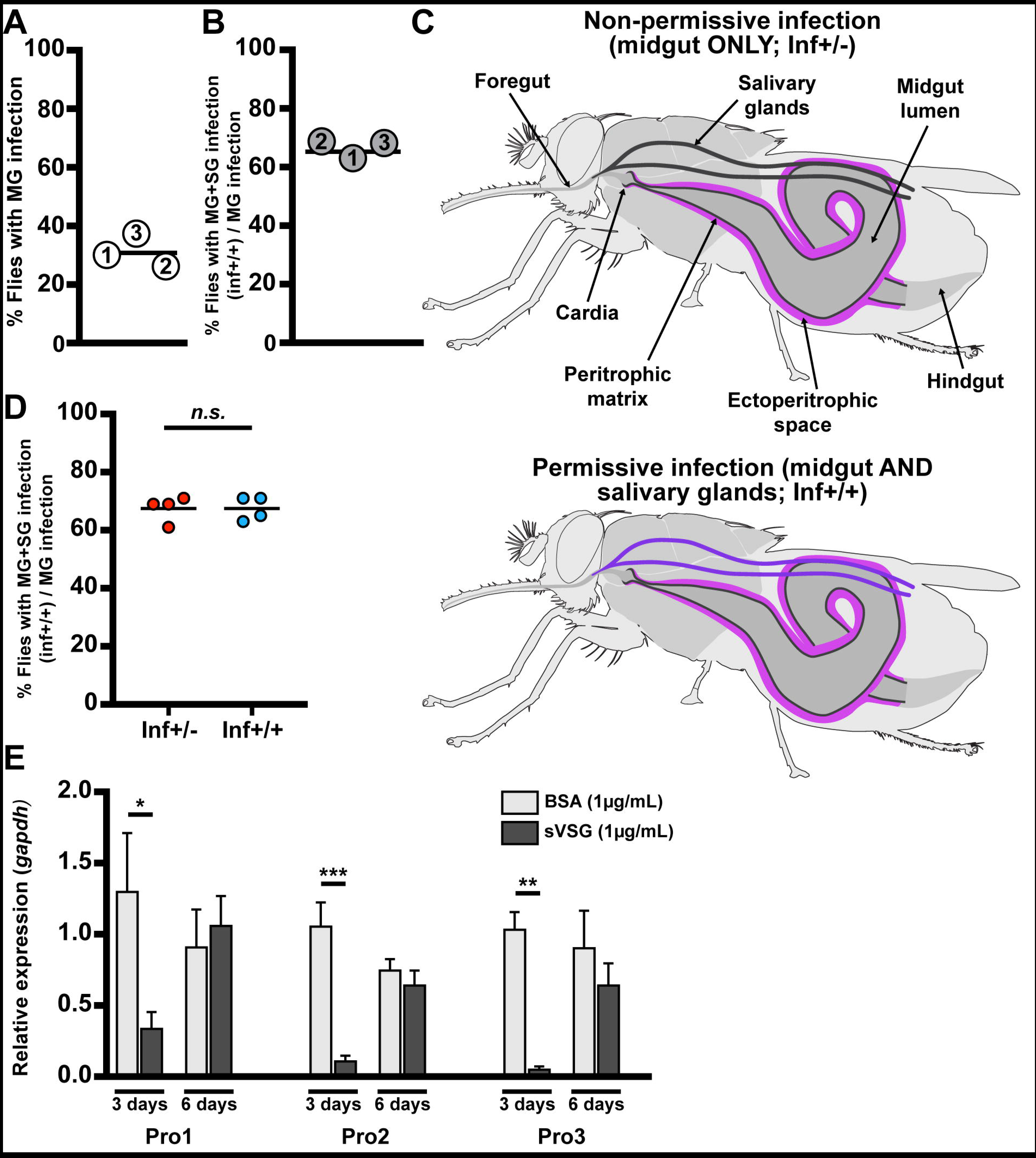
Dynamics of parasite infection in tsetse flies. (**A and B**) Midgut (MG) and salivary gland (SG) organs were dissected from 40-day old female flies subjected to a single parasite challenge as newly eclosed adults. (A) Percentage of flies harboring MG infections, and (B) percentage of MG infected flies that also presented SG infections. The numbers in each circle represent the three independent biological replicates. The black bars represent the mean of the three biological replicates (detailed numbers of flies used for the three biological replicates in S1 Dataset). (**C**) Schematic depiction of trypanosome localization in tsetse fly tissues. In the non-permissive flies (inf+/- state, shown in the upper scheme) only the midgut, including the cardia, is colonized by parasites, which reside in the ES (purple). In permissive flies (inf+/+ state, shown the lower scheme), parasites infect the fly’s and SGs (violet). (**D**) Percentage of permissive infections (inf+/+) following challenge at teneral stage with parasites obtained from the midgut of either inf+/- or inf+/+ individuals. Four independent experiments were performed for each treatment with 64 infected flies observed in total for each treatment. The black bar represents the mean of the four experiments. SG infection is independent of the initial inf+/- or inf+/+ status of the parasite used for fly infection (GLM, p=1, detailed model in S1 Dataset). (**E**) Expression of PM-associated genes *proventriculm-1, 2* and *3* (*pro1, 2, 3*) relative to the housekeeping gene *gapdh*, three and six days after flies received a blood meal supplemented with 1μ/mL of either BSA (control, light grey) or trypanosome-derived soluble variant surface glycoprotein (sVSG; dark grey). Each bar represents the average (± SEM) of five biological replicates. For each time point of each gene, a Student t-test was used to determine significant differences (* p<0.05; ** p<0.01; *** p<0.001).

We next investigated whether parasites residing in the non-permissive (inf+/-) gut infections suffer a developmental bottleneck that result in the selection of trypanosomes that are incapable of progressing towards metacyclic infections in the SG. We challenged two groups of teneral adults *per os* with *Trypanosoma brucei brucei* obtained from midguts of either inf+/+ or inf+/- flies. We observed a similar proportion of inf+/+ and inf+/- phenotypes regardless of the parasite population (inf+/+ or inf+/-) provided for the initial infection (Fig 1D). This indicates that trypanosomes in the inf+/- gut population are still developmentally competent, and can complete their cyclic development to SG metacyclics. Thus, we hypothesized that the cardia physiological environment may determine the developmental course of trypanosome infection dynamics.

Our prior studies on the role of the PM during the initial parasite colonization event showed that release of VSG from the ingested BSF parasites as they differentiate to PCF cells interferes with gene expression in the cardia resulting in loss of PM integrity early in the infection process. We thus checked to ensure that parasite re-entry into the gut lumen 6-10 dpa was not due to the residual effect of BSF-shed VSG on PM integrity. To do so we analyzed the expression of PM-associated genes (*pro1*, *pro2* and *pro3*) in cardia three and six days after supplementing flies with a bloodmeal containing purified VSG. Our results show that expression of the PM associated genes is significantly reduced at the day-three time point, but that their expression fully recovers by the day-six time point. These findings indicate that parasite re-entry into the gut lumen in the cardia is unlikely affected by loss of PM integrity that results from the initial VSG effects (Fig 1E).

### Parasites bypass the PM and enter into the gut lumen in permissive (inf+/+)cardia

To investigate the molecular aspects of the infection barriers preventing SG colonization that subsequently limit parasite transmission in the inf+/- group, we used the infection scheme described above and pooled infected cardia into inf+/- and inf+/+ groups (n=3 independent biological replicates per group, with ten cardia per replicate). For comparison, we similarly obtained dissected cardia from age-matched normal controls (called non-inf; n=3 independent biological replicates per group, with ten cardia per replicate). We next applied high-throughput RNA-sequencing (RNA-seq) to profile gene expression in the three groups of cardia. We obtained on average > 23M reads for each of the nine libraries, with 77.8% (non-inf), 75.4% (inf+/-) and 64.5% (inf+/+) of the total reads mapping to *Glossina morsitans morsitans* transcriptome (S1 Fig). The trypanosome reads corresponded to about 3.5% in non-permissive (inf+/-) dataset and about 11.9% in the permissive (inf+/+) dataset (Fig 2A). To estimate relative parasite densities in the two cardia infection states, we measured the expression of the trypanosome housekeeping gene *gapdh* in inf+/- and inf+/+ cardia by quantitative real time PCR (qRT-PCR) and normalized the values using tsetse *gapdh*. We noted significantly higher parasite gene expression values in the inf+/+ cardia compared to the inf+/- cardia samples (Student t-test, *p* = 0.0028; Fig 2B). We also confirmed that inf+/+ cardia had higher parasite density by microscopically counting trypanosome numbers in the dissected cardia organs using a hemocytometer (S2 Fig). Thus, the difference in the representative parasite transcriptome reads in the two infected groups of cardia is due to an increase in the number of trypanosomes residing in the inf+/+ cardia rather than an increase in parasite transcriptional activity. Interestingly, we noted no difference in the number of trypanosomes present in inf+/- and inf+/+ midguts (S2 Fig). Hence, it appears that parasite density either decreases in the cardia, or fewer parasites colonize the organ despite the fact that inf+/- and inf+/+ flies maintain similar parasite densities in their midguts.

**Fig 2.**
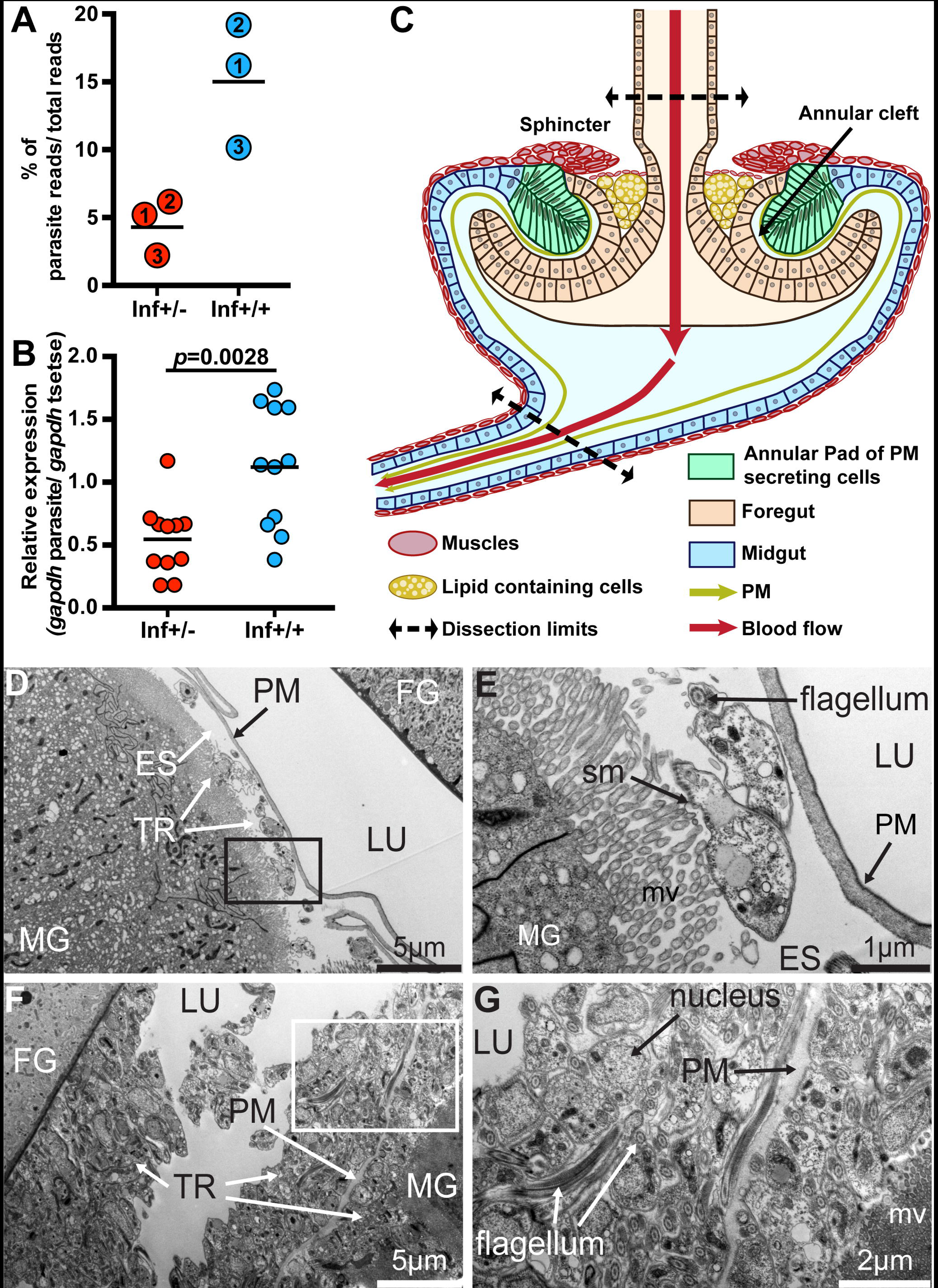
Trypanosome infection establishment process. (**A**) Percentage of RNAseq reads, relative to the total, that mapped to the parasite reference geneset from the three independent biological replicates. Inf+/- cardia is depicted in red circles and inf+/+ in blue circles. The numbers in circles represent the three biological replicates. The black bars indicate the mean of the three replicates (detailed numbers of flies used for the three biological replicates in S1 Dataset). (**B**) Abundance of trypanosome *gapdh* relative to tsetse *gapdh* determined from inf+/- cardia (shown by red circles) and inf+/+ cardia (shown by blue circles). The black bars indicate the mean of the replicates. The increase in relative abundance indicates an increase in parasite numbers in the inf+/+ cardia (Student t-test, *p=* 0.0028). (**C**) Schematic representation of the cardia organ based on microscopic observations. The cardia is composed of cells originating from the foregut (light orange) and midgut (light blue) at the junction of the foregut and midgut. Specialized midgut cells organized as an annular pad around the invaginated foregut secretes the PM (green) in the annular cleft formed between the foregut and midgut. Sphincter muscles that form a ring above the PM secreting cells, as well as the thin layer of muscle that surrounds large lipid-containing cells (shown in yellow), are indicated. The schematic indicates the upper and lower points where cardia were dissected for all experiments.In this schematic, the crop duct connecting the foregut prior to its invagination in the cardia is not presented. (**D-G**) Representative TEM micrographs showing cardia from inf+/- (D-E) and inf+/+ (F-G) individuals. (E) and (G) are magnified micrographs of the black and white boxes in (D) and (F), respectively. Cardia from six and five individuals from inf+/+ and inf+/- flies were imaged, respectively. MG: midgut; FG: foregut; ES: ectoperitrophic space; TR: trypanosomes; PM: peritrophic matrix; LU: lumen; mv: microvilli; sm: subcellular microtubules.

To detect the presence of parasites in the foregut, and to understand the different parasite developmental stages that could be present in the inf+/- and inf+/+ cardia, we investigated parasites by microscopy from these tissues at 40 dpa. We did not observe parasites in the foregut of inf+/- infections, confirming that they are are restricted from further development in the cardia in this group of flies. In the inf+/+ state however, we noted presence of many parasites in the foregut in all examined flies. We also looked at the relative presence of the different parasite developmental forms (short and long-epimastigote and trypomastigotes) populating the two different cardia phenotypes by examining parasite morphology and the localization of the nucleus and kinetoplast, as previously described [18, 19]. We observed that the majority of the parasites present in both cardia infection states on day 40 were trypomastigotes with fewer epimastigotes, and that no significant differences between the two cardia infection states was noted (S1 Dataset).

Tsetse’s cardia is composed of several different cell-types with potentially varying functions (schematically shown in Fig 2C; S3 Fig) [25–29]. These include an invagination of cells originating from the foregut, which are enclosed within an annular pad of columnar epithelial cells originating from the midgut. The cells occupying this pad secrete vacuoles that deliver components of the PM [27, 29]. The cardia organ is surrounded by muscles that form a sphincter around the foregut, which likely regulates blood flow during the feeding process. Additionally, large lipid-containing cells are localized under a layer of muscle below the sphincter. The function of these cells remains unclear. Microscopy analysis of infected cardia supported our previous molecular findings, as we observed fewer parasites in the cardia of inf+/- (Fig 2D; Fig 2E) when compared to inf+/+ flies (Fig 2F; Fig 2G). Parasites from the inf+/- cardia were restricted to the ES, whereas parasites were observed in both the ES and the lumen of inf+/+ cardia. Hence, the parasite populations resident in inf+/+ cardia had translocated from ES to the lumen, while parasites in inf+/- cardia failed to bypass the PM barrier. These data suggest that the cardia physiological environment may influence the parasite infection phenotype and transmission potential.

### PM is compromised in permissive (inf+/+) cardia but not in non-permissive (inf+/-) cardia

For succesful transmission to the next mammalian host, trypanosomes that reside in the ES of the midgut must traverse the PM barrier a second time to re-enter into the gut lumen, move forward through the foregut and mouthparts and colonize the SGs. Traversing the PM a second time is thought to occur near the cardia region [25, 29–31] due to newly synthesized PM likely providing a less robust barrier than in the midgut region. We investigated whether the functional integrity of the PM in the two different infection states varied in the cardia organ. We mined the non-inf cardia transcriptome dataset (S2 Dataset) and identified 14 transcripts associated with PM structure and function [6, 9, 32], which cumulatively accounted for 35.7% of the total number of reads based on CPM value (Fig 3A). The same set of genes represented 26.5% and 34.5% of the inf+/+ and inf+/- transcriptome data sets, respectively (Fig 3A). Thus, PM-associated transcripts are less abundant in the inf+/+ cardia relative to inf+/- and control cardia. We next evaluated the expression profile of PM-associated transcripts and identified those that are differentially expressed (DE) with a fold-change of ≥1.5 in at least one infection state compared to the control (non-inf) (Fig 3B). We observed a significant reduction in cardia transcripts encoding the major PM-associated proventriculin genes (*pro2*, *pro3*) in the cardia inf+/+, but not the cardia inf+/- dataset. Both Pro2 and Pro3 are proteinaceous components of the PM [6]. Interestingly, the expression of *chitinase* was induced in both inf+/- and inf+/+ datasets. Because Chitinase activity can degrade the chitin backbone of the PM, increased levels of its expression would enhance the ability of the parasites to bypass this barrier. Overall, the inf+/+ cardia expression profile we observed here is similar to the profile noted in the cardia 72 hours post BSF parasite acquisition early in the infection process [9]. Results from that study demonstrated that reduced expression of genes that encode prominent PM associated proteins compromised PM integrity, thus increasing the midgut parasite infection prevalence [9]. Loss of PM integrity in the inf+/+ state could similarly enhance the ability of parasites to traverse the PM to re-enter the gut lumen and invade the SGs.

**Fig 3.**
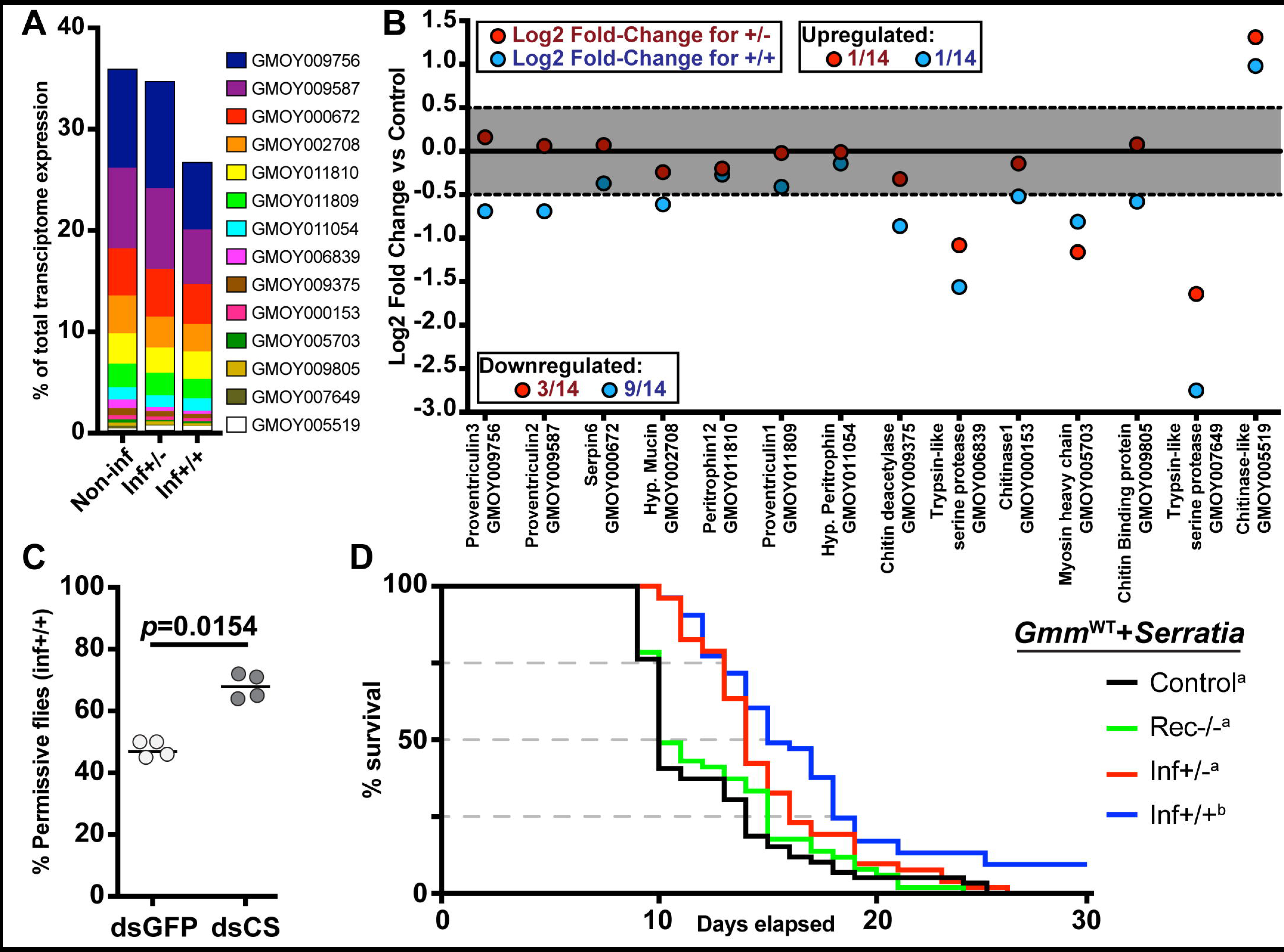
Parasite infection effects on PM synthesis in cardia inf+/+ and inf+/-. (**A**)
Percentage of transcripts that encode PM-associated proteins, relative to the total number of transcripts in each transcriptome. (**B**) Differential expression of PM-associated transcripts in cardia of flies that house inf+/+ (blue circles) and inf+/- (red circles) infections. Each dot represents the fold-change in expression of one transcript relative to the non-infected control. The grey area delineates fold-changes that are <1.5, and thus not statistically different from the control cardia (*p>* 0.05 after FDR correction). For each data point, the *Glossina* gene ID and function, based on BlastX annotation, is depicted on the x-axis. (**C**) SG infection prevalence in control (dsRNA-/p, dsGFP) and treatment (PM compromised; *dsRNA-chitin synthase*, dsCS) flies. The circles depict the percentage of flies that harbor both gut and SG infections. A total of four independent experiments were set-up for each group. The black bars indicate the mean of the four experiments. A total number of 66 and 63 infected flies were observed in the dsGFP and dsCS treatments, respectively. The dsCS treatment significantly increases trypanosome infection prevelance in tsetse’s salivary glands (GLM, Wald-test p=0.01408). Detailed counts and a complete statistical model are indicated in S1 Dataset. (**D**) Effect of cardia inf+/- and inf+/+ extracts on PM integrity. Survival of flies was monitored daily after *per os* treatment of 8 day-old flies with cardia extracts followed by *per os* treatment with *Serratia marcescens* 72 hours later. The Kaplan-Meyer curves show fly survival over time: cardia inf+/- (red), cardia inf+/+ (blue), or cardia from flies that recovered from infection (rec-/-, green) and cardia non-inf (black).Statistical analysis was performed using a full regression model followed by a pairwise test (details in S1 Dataset). Different letters next to fly group designations in the figure legend represent significantly different curves (p< 0.05).

### Reduction of PM integrity increases the prevalence of SG infections (inf+/+)

We hypothesized that PM integrity is a prominent factor in the ability of trypanosomes to traverse the barrier in the cardia and continue their migration to the SGs. We addressed this hypothesis by experimentally compromising the structural integrity of the PM in flies that harbored established gut parasite infections. We modified a dsRNA feeding procedure that targets tsetse *chitin synthase* (dsRNA-*cs*), which effectively inhibits the production of a structurally robust PM [7]. We challenged flies with BSF trypanosomes as teneral adults and then administered blood meals containing dsRNA-cs on day 6, 8 and 10 post parasite acquisition. This is the time interval when we expect the parasites colonizing the ES of the midgut to bypass the PM barrier in the cardia to re-enter into the lumen [19, 33, 34]. Control groups similarly received dsRNA targeting green fluorescent protein (dsRNA-gfp). Decreased expression of *chitin synthase* in the experimental dsRNA-cs group relative to the control dsRNA-gfp group was confirmed by qPCR analysis (S4 Fig). Twenty days post dsRNA treatment, midguts and SGs were microscopically dissected and the SG infection status scored. We detected SG infections in 68% of dsRNA-cs treatment group compared to 47% in dsRNA-gfp control group (Fig 3C). Thus, the PM compromised group of flies showed a significant increase in inf+/+ phenotype relative to the control group (GLM, Wald-test, p = 0.0154). These findings suggest that compromising the PM structure later in the infection process increases the proportion of gut infected flies that give rise to mature SG infections (inf+/+). Thus, tsetse’s PM acts as a barrier for parasite translocation from the ES to the lumen of the midgut, an essential step for successful SG colonization.

### Permissive (inf+/+) cardia extracts compromise PM integrity

We sought to determine if components of inf+/+ parasites infecting the cardia could manipulate cardia physiology to bypass the PM. For this, we used a modified version of a host microbial survival assay that was successfully used to evaluate PM structural integrity [7, 9, 35]. In this assay, tsetse with an intact PM fail to immunologically detect the presence of the entomopathogenic *Serratia marcescens* in the gut lumen. The bacteria thus proliferate uncontrolled in this environment, translocate into the hemocoel and kill the fly [7]. Conversely, when PM structure is compromised, the fly’s immune system can detect the presence of *Serratia* early during the infection process and express robust antimicrobial immunity that limits pathogen replication and increases host survival [7]. We provided mature adults blood meals supplemented with both entomopathogenic *Serratia* and heat-treated inf+/+ cardia extracts, while two age-matched control groups received either both *Serratia* and cardia extracts prepared from flies that had cleared their midgut infections (designated rec-/- for "recovered") or only *Serratia* (control). We found that survival of flies that received inf+/+ extracts was significantly higher than either of the two control groups (Fig 3D). These findings suggest that cardia inf+/+ extracts contain molecule(s) that negatively influence tsetse‘s PM integrity, thereby enabling these flies to more rapidly detect *Serratia* and express heightened immune responses to overcome this pathogen. Our transcriptional investigation indicated that PM associated gene expression decreased in the inf+/+ state but not in the inf+/- state (Fig 3B). In the survival assay we described above, we fed flies with cardia extracts from inf+/- containing heat-killed parasites at an equivalent quantity as the one used in the inf+/+ state. When supplemented with the cardia+/- extracts, the survival of flies was decreased to the same level as the two controls, suggesting that extract from cardia inf+/- did not compromise PM integrity (Fig 3D). Collectively, these findings confirm that the parasites in cardia inf+/- differ in their ability to interfere with PM integrity when compared with those in the cardia inf+/+ state. This suggests that parasites in inf+/+ cardia display a different molecular dialogue with tsetse vector tissues.

### Only parasites from cardia inf+/+ bypass the PM barrier

To understand the cardia-trypanosome interactions, we investigated the parasite populations in the inf+/+ state by transmission electron microscopy (TEM) analysis. We observed that trypanosomes aggregate in the annular cleft formed between the foregut and the midgut parts of the cardia where PM components are synthesized (Fig 4A; S5 Fig). Tsetse’s PM is composed of three layers; a thin layer that is electron-dense when observed with TEM, a thick layer that is electron-lucent when observed with TEM and a third layer that is not distinguishable when observed with TEM [36]. The newly synthesized PM in the annular cleft is formed by secretions from the annular pad of epithelial cells [27, 29], hence lacking the typical electron-dense and electron-lucent layers observed in the fully formed PM in the midgut (Fig 4B; Fig 4C). From the six inf+/+ cardia analyzed by TEM, we observed trypanosomes embedded in the newly secreted PM as well as present in the lumen. In fact, we had shown above that the expression of putative PM-components decreased in cardia inf+/+ (Fig 3B), and the structural integrity of the PM is compromised based on the *Serratia* detection assay (Fig 3D). Thus, the structurally weakened PM could enable the trypanosomes to bypass this barrier in inf+/+ flies. Our EM observations (all six inf+/+ cardia) also showed that parasites assemble into compact masses (similar to the previously reported "cyst-like" bodies [33]) in between the layers of the PM (Fig 4C; S6 Fig). In three of the six infected inf+/+ cardia analyzed, we noted that the electron-dense layer of the PM restricting a cyst-like body appeared disrupted, which could enable the entrapped parasites to escape the barrier (S7 Fig).

**Fig 4.**
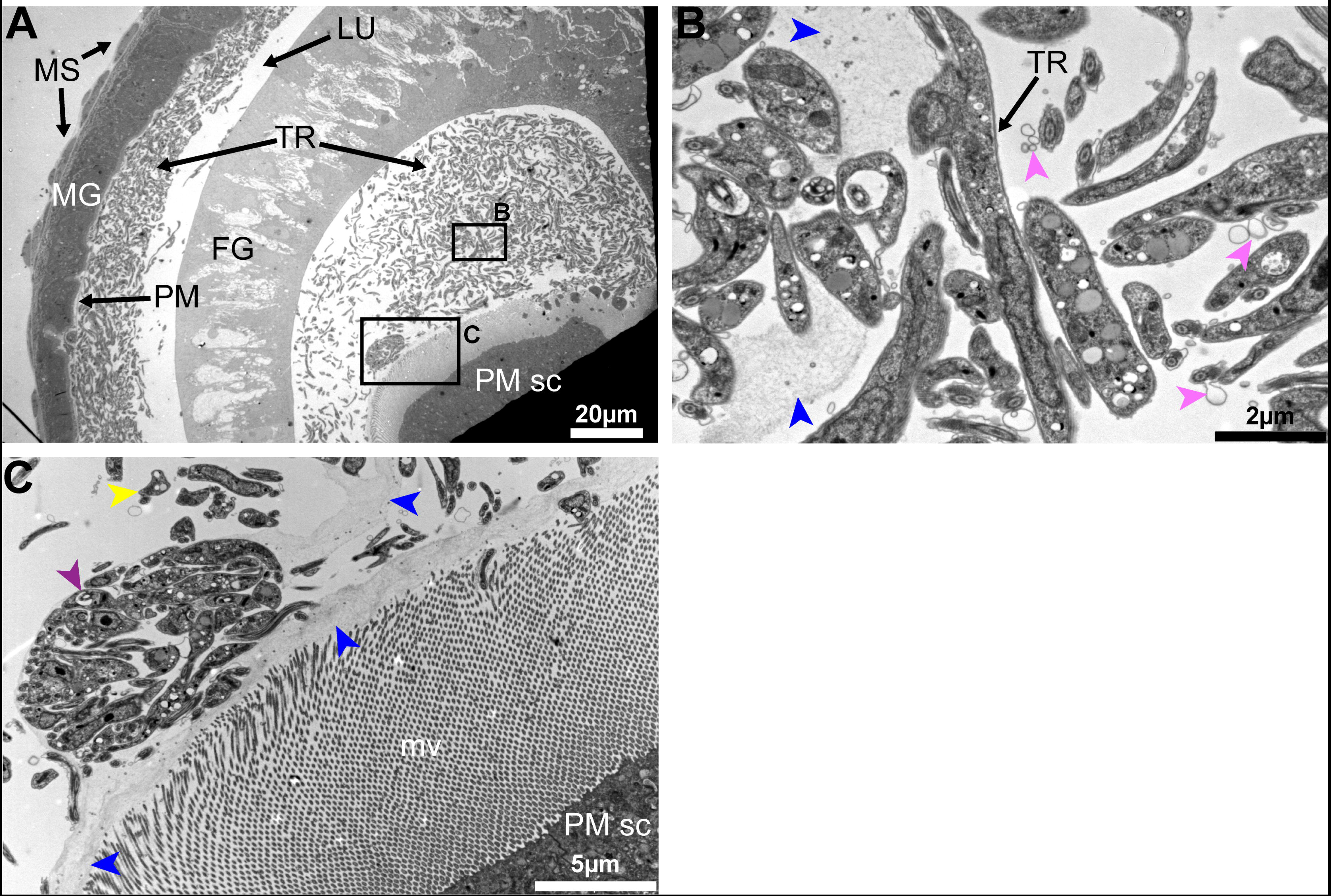
Trypanosome-PM interactions in cardia from inf+/+ tsetse. (**A-D**) Ultrastructure of PM secreting cells in the cardia from inf+/+ flies. (A) Trypanosomes are observed in mass in the lumen. (B-C) Magnified micrographs of the black boxes shown in (A).(B)
Trypanosomes are observed embedded in the secreted matrix (blue arrowheads). (C) In this niche parasites are observed in cyst-like bodies (purple arrowhead), and can also be observed out of the PM secretions (yellow arrowheads). Parasite secreted extracellular vesicles are observed (pink arrowheads). Micrographs in this image represent one of six biological replicates analyzed. MG: midgut; FG: foregut; TR: trypanosomes; MS: muscles; PM: peritrophic matrix; PM sc: PM secreting cells; mv: microvilli.

The parasite aggregates we observed in the cardia near the site of PM secretion could represent a social behavior that influences cardia-trypanosome interactions and ultimately parasite transmission success. *In vitro*, trypanosomes are capable of displaying a similar social behavior termed ‘social motility’ (SoMo) [37]. In this situation early-stage PCF parasites (similar to the forms that colonize the fly midgut) cluster and migrate together on semi-agar plates [38]. In the tsetse vector, phases where trypanosomes group in clusters and move in synchrony have been observed during the infection process independent of the developmental stage of the parasite [34]. Furthermore, parasites co-localize in the cardia near the cells that produce the PM [34], similar to our EM observations. By forming aggregates, trypanosomes could enhance their ability to resist adverse host immune responses and/or escape the ES by crossing through the newly secreted layers of the PM. In addition, the parasites can also actively compromise the PM integrity at this site, as suggested by the PM integrity assay (Fig 3D), but the parasite components that interfere with host functions as such remain to be determined. We also observed extracellular vesicles associated with trypanosomes in TEM images, which could potentially carry molecules that interact with host cells or PM structure (Fig 4).

To understand the parasite-PM interactions in the cardia inf+/- state, we similarly investigated the parasite populations residing in the cardia inf+/- samples by TEM analysis. We observed that parasites in cardia inf+/- are not present in the lumen and are thus unable to escape the ES where the newly synthesized PM is secreted (Fig 5). We noted that high densities of parasites are either lining along the PM secreting cells (Fig 5A; Fig 5B) or are embedded in the PM secretions (Fig 5C; Fig 5D). Typanosomes observed in this region also presented multiple vacuolation and nuclear condensation, which are indicative of cell death processes in these parasites. Contrary to the cardia inf+/+ transcriptome data, the expression of the majority of PM-associated genes in cardia inf+/- are not significantly decreased (Fig 3B). Moreover, the *Serratia* assay we applied by co-feeding flies cardia inf+/- extracts indicated no compromise of PM integrity as these group of flies did not survive the bacterial infection (Fig 3D). Thus, it appears that parasites in the cardia inf+/- are restricted by the PM to remain in the ES even at its point of secretion. Also, while cyst-like bodies were frequent in the cardia inf+/+, only a few cyst-like bodies could be observed in cardia inf+/-. Finally, the presence of many physiologically unfit trypanosomes indicates that the inf+/- state represents a hostile environment for the parasite, restricting its survival and transmission (Fig 5C; Fig 5D).

**Fig 5.**
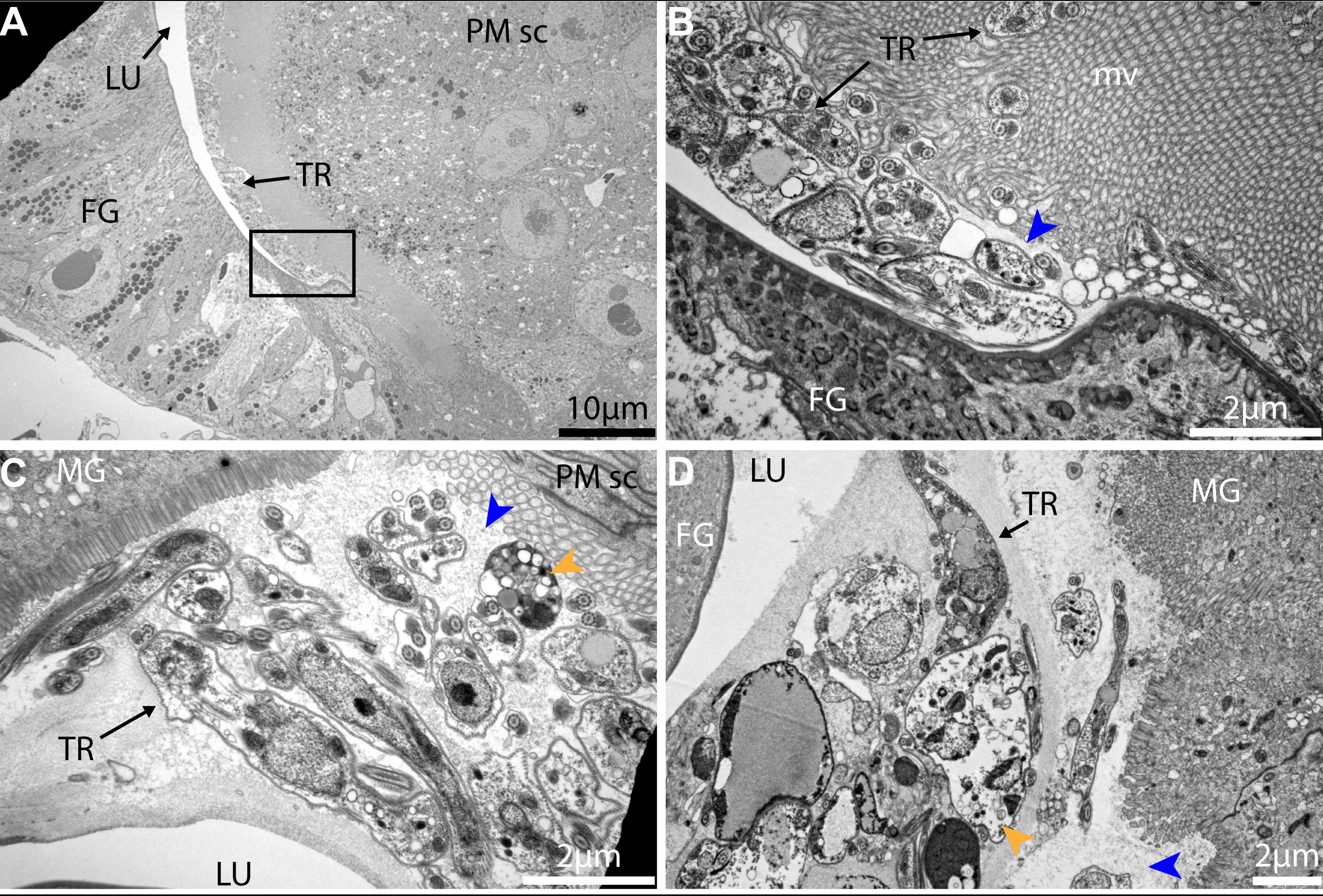
Trypanosome-PM interactions in cardia from inf+/- tsetse. (A-D) Ultrastructure of cardia inf+/- near the PM secreting cells. (B) is a magnification of the black frame in (A) showing the PM (blue arrowhead). (C) and (D) two independent cardia organs showing the same region near PM secreting cells. Trypanosomes are observed packed within the ES near the location of PM secretion. At this point, several trypanosomes observed present vacuolation and nuclear condensation (orange arrowheads) indicative of cell death. Micrographs in this image represent three of five biological replicates analyzed. MG: midgut; FG: foregut; TR: trypanosomes; PM: peritrophic matrix; PM sc: PM secreting cells; mv: microvilli.

### The cardia from non-permissive infections (inf+/-) is dysfunctional at the cellular level

To understand the factors that can successfully inhibit parasite survival in the cardia inf +/-, we examined the inf+/- and inf+/+ cardia datasets relative to the control non-inf state for differential vector responses. We found that 25% (2093) of the total transcripts identifed were differentially expressed (DE). Of the DE transcripts, 31% (646) were shared between the two cardia infection phenotypes, while 36% (756) and 33% (691) were unique to the inf +/- and inf+/+ infection phenotype, respectively (S8 Fig). Of the shared DE transcripts, 89% (576) were similarly regulated between the inf+/+ and inf+/- states while 11% (70) were uniquely regulated in the two infection states. For putative functional significance, we selected transcripts presenting a fold-change of ≥2 between any comparison and a mean CPM value of ≥50 in at least one of the three cardia states. We identified 576 transcripts that were modified in the presence of trypanosomes independent of the cardia infection phenotype, hence representing the core response of the cardia against the parasite infection (S8 Fig). Among these core responses were three antimicrobial peptide (AMP) encoding genes, including two *cecropins* with fold-changes of >200 (GMOY011562) and >280 (GMOY011563), and *attacin D*, with a fold change of >27. Production of AMPs by midgut epithelia is among the major anti-trypanocidal responses, and the fact that both cardia inf+/+ and cardia inf+/- expressed these genes at the same level indicates that the ability of inf+/- flies to restrict trypanosomes in the ES is unlikely driven by an AMP-related immune response. We next investigated DE transcripts unique to the two infection phenotypes (S8 Fig). Two putative immunity products, Immune responsive product FB49 and *serpin 1*, were expressed 223 and 74 times higher, respectively, in inf+/+ compared to inf +/ cardia. Both of these products are induced upon microbial challenge in the tsetse [13, 39]. Additionally, *ferritin* transcript abundance was >2 times higher in inf+/+ compared to inf +/ cardia. In the subset of transcripts specifically more abundant in the cardia inf+/-, we noted two transcripts encoding proteins involved in the circadian clock, Takeout and Circadian clock-controlled protein, which were 600 and 2 times more abundant relative to cardia non-inf, respectively. Also, transcripts encoding Kazal-type 1 protein, a protease inhibitor, and Lysozyme were more abundant in cardia inf+/-. Given that no single immune-related gene product could explain the cardia inf+/- ability to restrict trypanosomes in the ES, we chose to further evaluate the cardia cellular physiology under the inf+/+ and inf+/- infection states.

To obtain a global snapshot of cardia functions that could influence parasite infection outcomes, the DE cardia transcripts between control (non-inf) and either cardia inf+/- or cardia inf+/+ datasets were subjected to Gene Ontology (GO) analysis (using Blast2GO) (S3 Dataset). We noted 88 GO terms that were significantly down-regulated preferentially in the inf+/- state, while only 15 GO terms were significantly down-regulated in the inf+/+ state. The 88 GO terms detected in the inf+/- dataset included 5, 11 and 67 terms associated with mitochondria, muscles and energy metabolism, respectively.

To understand the physiological implications of the inf+/- infection phenotype in the cardia, we investigated the transcriptional response of the organ as well as the ultrastructural integrity of the mitochondria and muscle tissue. Gene expression patterns indicate that mitochondrial functions are significantly down-regulated in the inf+/- cardia relative to the inf+/+ state (Fig 6A). More specifically, the putative proteins associated with energy metabolism, including the cytochrome c complex, the NADH-ubiquinone oxidoreductase and the ATP-synthase that function at the organelle’s inner membrane, were very reduced. Loss of mitochondrial integrity was further demonstrated by microscopic analysis of cardia muscle cells (Fig 6B-D; S9 Fig) and fat-containing cells (Fig 6E-G). In the cardia inf+/- phenotype, TEM observations showed mitochondrial degradation around myofibrils associated with muscle cells (Fig 6C), while few such patterns were noted in the control cardia (Fig 6B) and cardia inf+/+ (Fig 6D). The mitochondria within the lipid containing cells of both inf+/- and inf+/+ presented a disruption in the organization of their cristae, suggesting a disruption of the inner membrane (Fig 6F-G), in support of transcriptomic level findings (Fig 6A).

**Figure 6.**
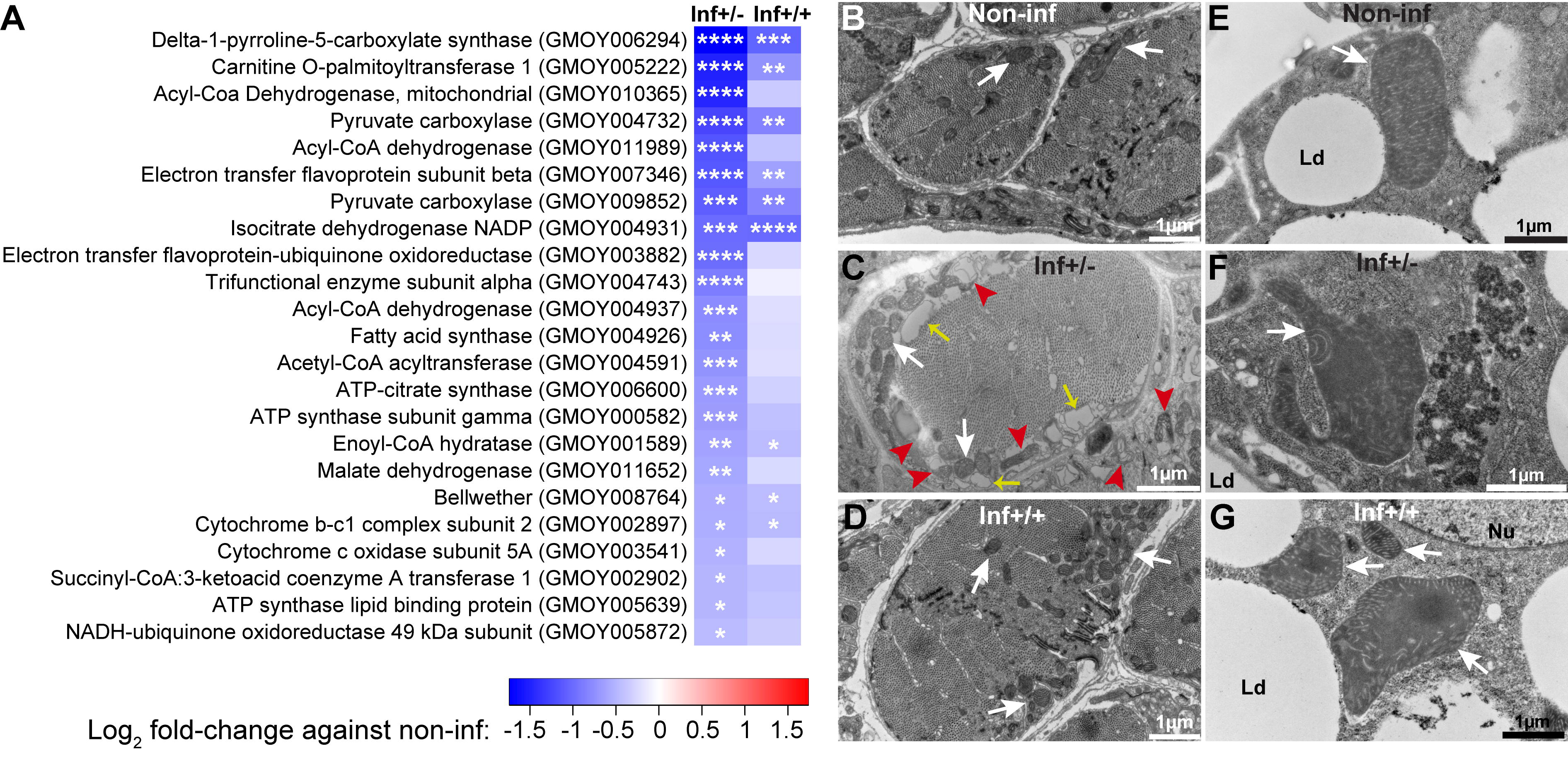
Mitochondrial integrity in cardia from inf+/+ and inf+/-tsetse. (**A**) Effect of infection on mitochondria related gene expression. Heatmap generated from the fold-changes between control and either inf+/- or inf+/+ cardia. The * denote the level of significance associated with the DE of specific transcripts (*p<0.05; **p<0.01; ***p<0.001; ****p<0.0001). (**B-D**) Ultrastructure of the sphincter myofibrils in control non-inf (B), inf+/-(C) and inf+/+ (D) cardia. White arrows show the mitochondria, red arrowheads show patterns of mitochondria degradation, and yellow arrows show dilatation of sarcoplasmic reticulum. (**E-G**) Ultrastructure of giant lipid-containing cells in control non-inf (E), inf+/- (F) and inf+/+ (G) cardia. In both infection phenotypes, mitochondria cristae appear disogarnized compared to control. Micrographs in this image represent one of three, five and six of biological replicates from cardia non-inf, inf+/- and inf+/+, respectively. White arrows show the mitochondria. Ld, lipid droplets; Nu, nucleus.

In addition to putative mitochondrial proteins, we found that the expression of genes encoding structural proteins responsible for muscle contraction, such as myosin and troponin, is also significantly reduced upon infection, particularly in the cardia inf+/- state (Fig 7A). Electron microscopy analysis also revealed a disorganization of the Z band of sarcomeres in muscle tissue surrounding the midgut epithelia in inf+/- cardia, but not in the control and inf+/+ cardia (Fig 7B-D). Extensive loss of muscle integrity was noted along the midgut epithelia in the inf+/- state. In addition, dilatation of the sarcoplasmic reticulum, muscle mitochondria swelling and vacuolation were observed, suggesting compromised muscle functions associated with this infection phenotype (S9 Fig). The detrimental effects of trypanosome infection on cardia structure and function are more apparent in the inf+/- compared to inf+/+ state, despite the higher number of parasites present during the latter phenotype.

**Fig 7.**
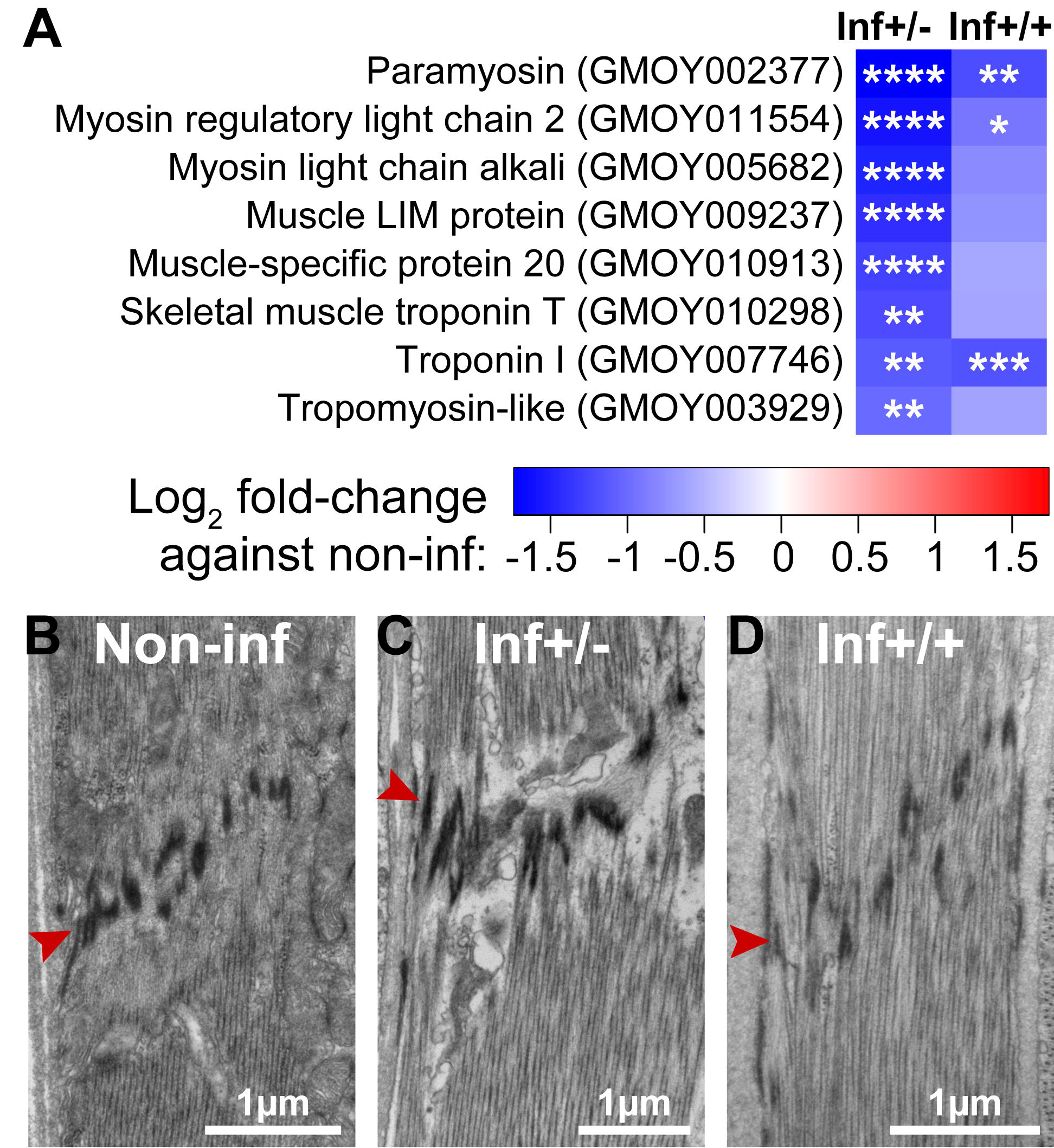
Muscle structural integrity in cardia from inf+/+ and inf+/-. (**A**) Effect of infection on cardia muscle related gene expression. Heatmap generated from the fold-changes between control and either inf+/- or inf+/+ cardia. The * denote the level of significance associated with the DE of specific transcripts (*p<0.05; **p<0.01; ***p<0.001; ****p<0.0001). (**B-D**) Ultrastructure of a sarcomere from muscles surrounding non-inf (B), inf+/- (C) and inf+/+ (D) cardia. The red arrowhead indicates the Z band structure associated with sarcomeres. Micrographs in this image represent one of three, five and six of biological replicates from cardia non-inf, inf+/- and inf+/+, respectively.

### Oxidative stress restricts parasite infections in cardia inf+/-

Mitochondria produce reactive oxygen intermediates (ROIs) [40], which in excess can damage the organelle and surrounding cellular structures [41, 42]. The structural damage we observed in mitochondria, muscle tissue and fat cells of inf+/- cardia is symptomatic of oxidative stress [43]. Additionally, our TEM observations demonstrate that parasites in inf+/- cardia exhibit cell-death patterns such as vacuolation and swelling (Fig 8A; Fig 8B), while parasites in inf+/+ cardia appear structurally intact (Fig 2F; Fig 4). Because ROIs modulates trypanosome infection outcomes in tsetse [15, 44], we hypothesized that ROIs may be responsible for controlling trypanosomes in inf +/- cardia and for producing an oxidative environment that concurrently results in tissue damage. We observed a significant increase of peroxide concentrations in both inf+/- (406nM; TukeyHSD posthoc test, p<0.0001) and inf+/+ (167nM; TukeyHSD posthoc test, p=0.0008) cardia relative to the control cardia (19 nM), with peroxide levels significantly higher in the inf+/- state (TukeyHSD posthoc test, p<0.0001) (Fig 8C). When we experimentally decreased oxidative stress levels in infected flies by supplementing their blood meal with the antioxidant cysteine (10μM) (Fig 8D), 85% of midgut infected flies developed SG infections, while only 45% of midgut infected flies had SG infections in the absence of the antioxidant (GLM, Wald-test p<0.001). Our results indicate that the significantly higher levels of ROIs produced in the inf+/- cardia may restrict parasite infections at this crucial junction, while lower levels of ROIs present in the inf+/+ cardia may regulate the parasite density without impeding infection maintenance.

**Fig 8.**
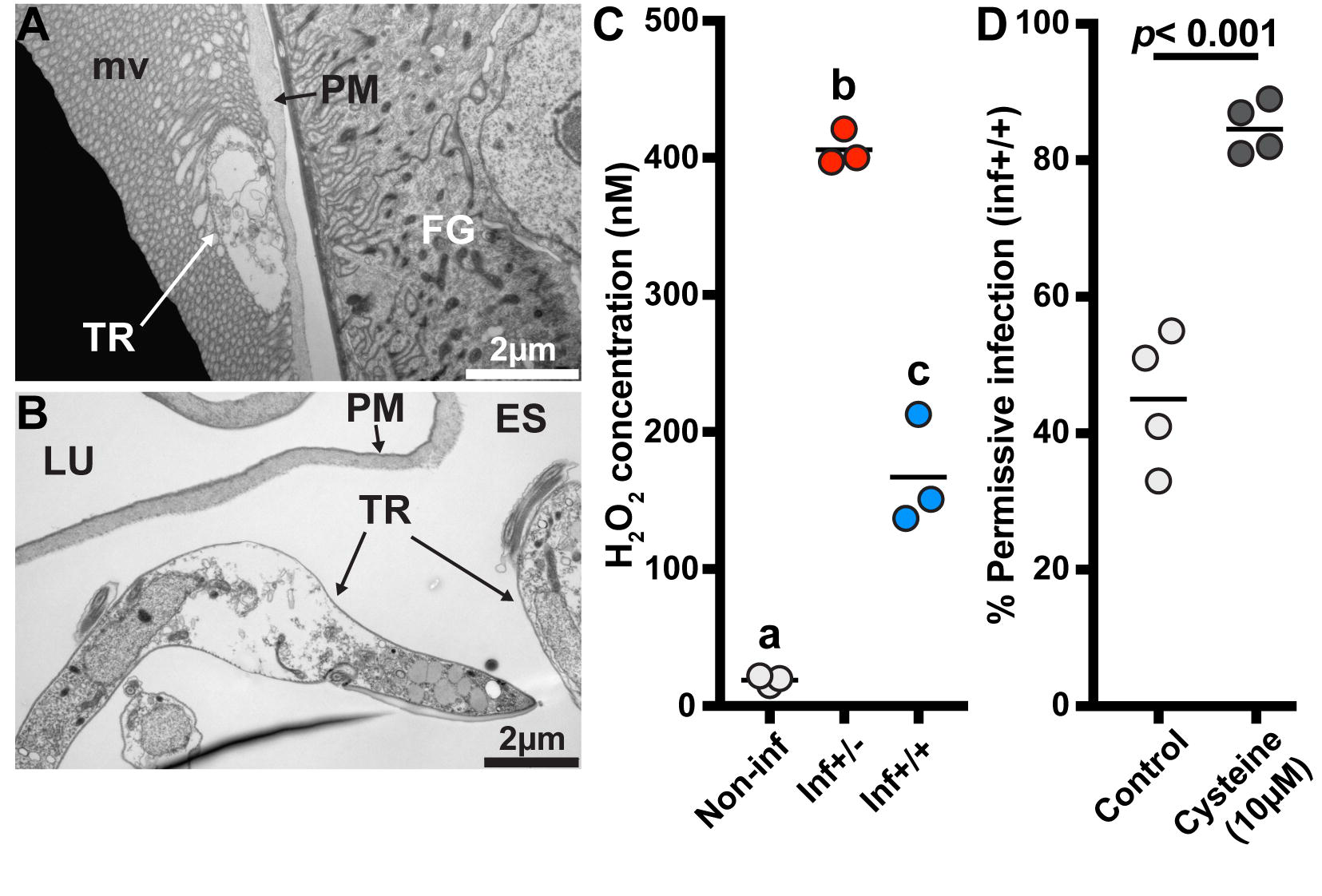
Influence of oxidative stress on infection status. (**A-B**) Electron microscopy observations of trypanosomes presenting cell-death swelling within the ES of cardia from inf+/- flies. Micrographs in this image represent two of five of biological replicates. FG: foregut; TR: trypanosomes; PM: peritrophic matrix; LU: lumen. (**C**) Comparison of peroxide levels in cardia from non-infected (white circles), inf+/- (red circles) and inf+/+ (blue circles) flies 72 hours after a blood meal. Each dot represents an independent quantification of 10 pooled cardia. The black bars indicate the mean of the 3 replicates. Statistical analysis was conducted using a one-way ANOVA followed by a TukeyHSD posthoc test for pairwise comparisons. Statistical significance is represented by letters above each condition, with different letters indicating distinct statistical groups (p< 0.05). (**D**) SG infection prevalence in normal and anti-oxidant (cysteine) treated flies. The circles depict the percentage of flies that harbor both gut and SG infections in the cysteine treated (10μM) and control groups. A total of 4 independent groups were set-up for each treatment (91 and 89 infected flies were observed in the control and cysteine treatment groups, respectively). The black bars indicate the mean of the 4 experiments.Cysteine treatment significantly increases trypanosome infection prevelance in tsetse’s salivary glands (GLM, Wald-test p<0.001). Detailed counting data and complete statistical model are indicated in S1 Dataset.

Homeostasis of redox balance is one of the most critical factors affecting host survival during continuous host-microbe interaction in the gastrointestinal tract [45]. In the mosquito *Anopheles gambiae*, increased mortality is observed when ROIs are produced in response to *Plasmodium berghei* infections [46]. A similar trade-off expressed in the inf+/- cardia may restrict parasite infections while causing collateral damage to essential physiologies. Conversely, strong anti-parasite responses that compromise essential physiologies are absent in the cardia of the inf+/+ group, thus allowing the parasites to continue their journey to colonize the SG and successfully transmit to a new host. Additionally, flies with SG parasite infections also suffer from longer feeding times due to suppressed anti-coagulation activity in the SG, which may further help with parasite transmission in this group of flies [47].

## Conclusion

Trypanosome transmission by tsetse reflects a tug-of-war that begins with parasite colonization of the midgut and ends when parasites are transmitted to the next vertebrate via saliva. Initially during the infection process, BSF trypanosome products manipulate tsetse vector physiology to bypass the gut PM to colonize the midgut ES [9]. Our results show that to successfully colonize the SG, trypanosomes may again manipulate tsetse physiology to escape the midgut ES for access to the foregut, and subsequently to the SG. To re-enter the lumen, it is hypothesized that trypanosomes cross the PM in the cardia where newly synthesized PM is less structurally organized and hence can provide an easy bypass [25, 29–31]. Here, we provide evidence in support of this hypothesis by showing that in flies where trypanosomes successfully colonize the SG, the parasites are accumulating in the region where the PM is newly secreted, and are observed both embedded in the PM secretions and free in the lumen (summarized in Fig 9). To facilitate their passage, components of trypanosomes in the cardia can apparently manipulate PM integrity by influencing the expression of PM-associated genes through molecular interference, the mechanisms of which remain to be studied. Alternatively, trypanosome-produced molecules may directly reduce the integrity of the PM as a barrier.

**Fig 9.**
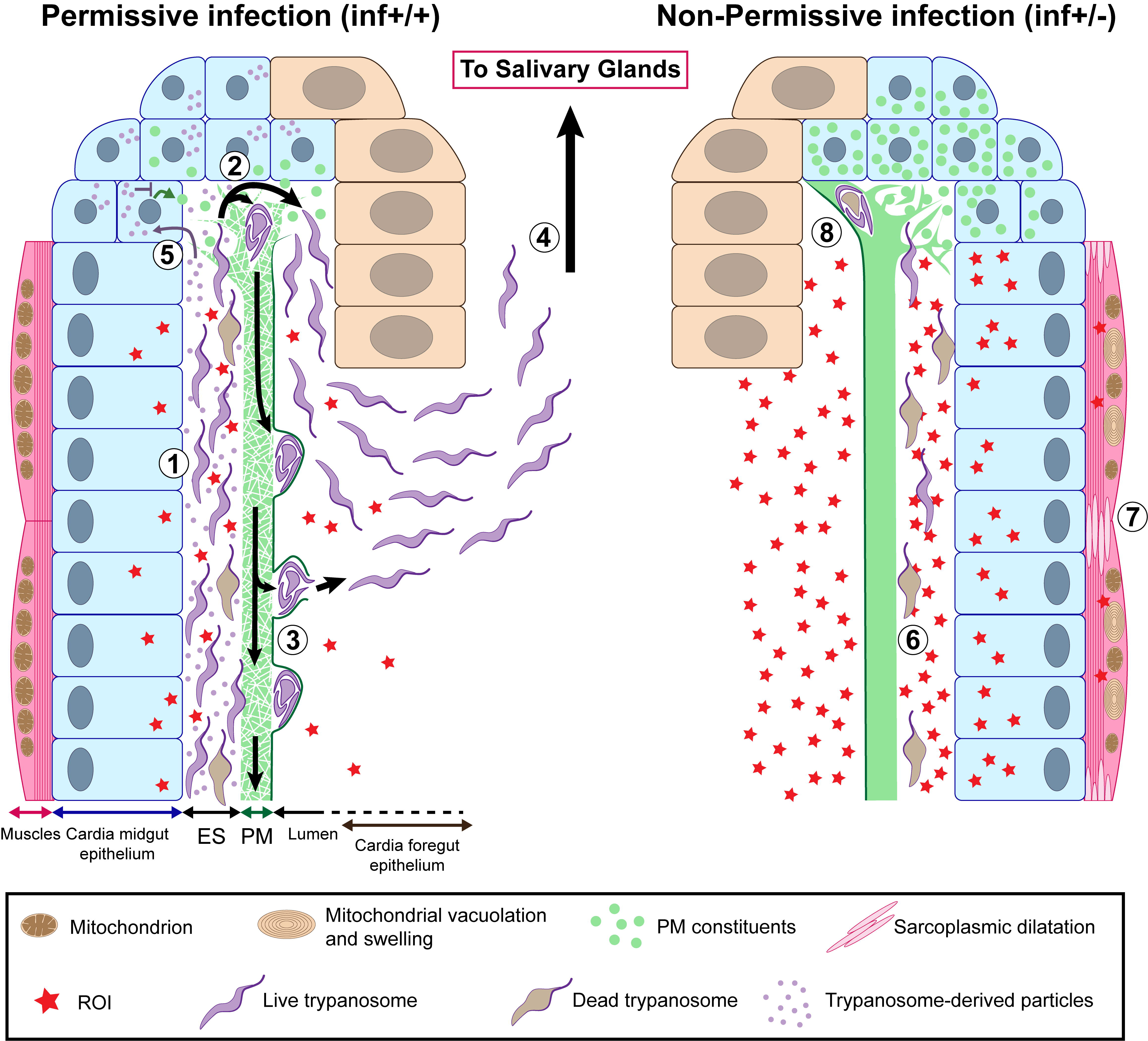
Model illustrating permissive (inf+/+) and non-permissive (inf+/-) infection phenotypes in tsetse’s cardia. African trypanosomes must pass through the tsetse vector in order to complete their lifecycle and infect a new vertebrate host. Following successful infection of tsetse’s midgut, *Trypanosoma brucei* parasites either remain trapped indefinitely in this environment (non-permissive infection, inf +/-) or migrate to the fly’s SG where they subsequently transmit to a vertebrate host (permissive infection, inf +/+). For a inf+/+ infection to occur, trypanosomes must successfully circumvent several immunological barriers, including the cardia-synthesized PM. In this situation, parasites that have accumulated in the ES (1) of the cardia traverse the structurally compromised PM at its site of synthesis (where the matrix is most diffuse and fragile) (2) (Fig 4; S5 Fig). The cyst-like bodies of parasites are observed within layers of the PM (S6 Fig), at which point they may force their way out by breaking through the structure’s electron-dense layer (S7 Fig). Otherwise they may stay enclosed within and move along the gut with the continuously generated PM (3) [33]. Parasites that have successfully translocated to the cardia lumen then migrate to the foregut and salivary glands (4). In an effort to facilitate their passage of the PM, trypanosomes may interfere with PM synthesis by secreting modulatory molecules (Fig 3D; Fig 4B) that are taken up by PM synthesizing cells (5). These molecules may subsequently inhibit expression of genes that encode proteinaceous components (Pro1, Pro3, etc) of the matrix (Fig 3B) and trypanocidal reactive oxygen intermediates (ROIs). In non-permissive infections, a relatively small number of parasites reach tsetse’s cardia (Fig 2A; Fig2B; S2 Fig), but they appear damaged or dead (6) (Fig 8A; Fig 8B). The inability for the parasite to sustain in the inf +/- cardia environment is likely caused by a relatively high concentration of ROIs in this environment (Fig 8C). ROI-mediated regulation of the parasite population comes with collateral damage to cardia tissues (Fig 6), especially the muscles lining the outer border of the organ, which present sarcoplasmic dilatation and mitochondrial vacuolation and swelling (7) (Fig 7; S9 Fig). In the inf +/- cardia, PM synthesis is not affected (Fig 3B), probably due to the absence of trypanosome-derived molecules interfering with the PM production (Fig 3D). Ability to restrict parasites in the cardia prohibits the trypanosomes from translocating to the cardia lumen for subsequent transmission (8).

The presence of trypanosomes in the cardia triggers immune responses which include the production of ROIs. In flies where midgut infections fail to reach the SG (inf+/-), increased levels of peroxide produced in the cardia may restrict parasite survival and prevent them from further development in the fly. Given that the inf+/- phenotype is costly and leads to collateral damage in the cardia tissues of infected flies, it is possible that flies may be able to sustain this phenotype under laboratory conditions where resources are abundant for a minimal effort. It remains to be seen if the inf+/- phenotype could sustain in natural populations in the field. Because in field infection surveillance studies estimating the time of initial parasite infection acquisition is not possible, concluding the cardia infection status in natural populations is difficult. It may however be possible to initiate parasite infection experiments using field-caught teneral flies to partially evaluate the potential colony-bias that could arise under insectary conditions using fly lines that have been kept in captivity for many years.

Trypanosome colonization of tsetse’s SG could represent a trade-off where vector tolerance to parasites leads to minimal self-inflicted collateral damage. Interestingly, different tsetse species may have evolved varying strategies to defend against parasitism. For instance, under similar laboratory conditions and using the same parasite strain for infection, *Glossina pallidipes* heavily defends against the initial infection, as the occurrence of the inf+/- phenotype in this species is rare despite similar resistance to SG transmission [48]. On the other hand, the closely related species *G. morsitans*, which we investigate here, has developed a different strategy to combat against parasite transmission [48]. *Glossina morsitans* presents a less efficient defense against the initial parasite infection in the midgut compared to *G. pallidipes*, but can similarly control the parasite transmission by restricting SG infections in midgut infected flies. Investigating the causes leading to this drift in strategies could lead to the development of new control strategies based on enhancing the immune defenses of the vector against parasites. Our work highlights the central role tsetse’s PM plays in parasite-vector interactions and infection outcome. This work opens up the possibility for exploiting this matrix as a target for vector control strategies to enhance its barrier function to block parasite transmission.

## Methods

### Ethical consideration

This work was carried out in strict accordance with the recommendations in the Office of Laboratory Animal Welfare at the National Institutes of Health and the Yale University Institutional Animal Care and Use Committee. The experimental protocol was reviewed and approved by the Yale University Institutional Animal Care and Use Committee (Protocol 2014-07266 renewed on May 2017).

### Biological material

*Glossina morsitans morsitans* were maintained in Yale’s insectary at 24°C with 50-55% relative humidity. All flies received defibrinated bovine blood (Hemostat Laboratories) every 48 hours through an artificial membrane feeding system. Only female flies were used in this study. Bloodstream form *Trypanosoma brucei brucei* (RUMP 503) were expanded in rats. Flies were infected by supplementing the first blood meal of newly eclosed flies (teneral) with 5×10^6^ parasites/ml. Where mentioned, cysteine (10μM) was added to the infective blood meal to increase the infection prevalence [15].

For survival assays, *Serratia marcescens* strain Db11 was grown overnight in LB medium. Prior to supplementation with *Serratia*, the blood was inactivated by heat treatment at 56°C for 1 hour as described in [7].

### mRNA Library Constructions and Sequencing

At day 40 post parasite challenge, all flies were dissected 48 hours after their last blood meal, and midgut and salivary glands (SG) were microscopically examined for infection status. Flies were classified as inf+/+ when infection was positive in both the midgut and the SG, as inf+/- when infection was positive in the midgut but negative in the SG. Cardia from inf+/+ and inf+/- flies were dissected and immediately placed in ice-cold TRIzol (Invitrogen). For each infected group, inf+/+ and inf+/-, 10 cardia were pooled into one biological replicate and three independent biological replicates were obtained and stored at −80°C prior to RNA extraction. Similarly, three independent biological replicates containing 10 cardia from age-matched flies that had only received normal blood meals (non-inf) were prepared. Total RNA was extracted from the nine biological replicates using the Direct-zol RNA Minipreps kit (Zymo Research) following the manufacturer instructions, then subjected to DNase treatment using the Ambion TURBO DNA-free kit AM1907 (Thermo Fisher Scientific). RNA quality was analyzed using the Agilent 2100 Bioanalyzer RNA Nano chip. mRNA libraries were prepared using the NEBNext Ultra RNA Library Prep Kit for Illumina (New England BioLabs) following the manufacturer recommendations. The nine libraries were sequenced (single-end) at the Yale Center for Genome Analysis (YCGA) using the HiSeq2500 system (Illumina). Read files have been deposited in the NCBI BioProject database (ID# PRJNA358388).

### RNA-seq data processing

Using CLC Genomics Workbench 8 (Qiagen), transcriptome reads were first trimmed and filtered to remove ambiguous nucleotides and low-quality sequences. The remaining reads were mapped to *Glossina morsitans morsitans* reference transcriptome GmorY1.5 (VectorBase.org). Reads aligning uniquely to *Glossina* transcripts were used to calculate differential gene expression using EdgeR package in R software [49].

Significance was determined using EdgeR exact test for the negative binomial distribution, corrected with a False Discovery Rate (FDR) at P<0.05.

Identified genes were functionally annotated by BlastX, with an E-value cut-off of 1e^-10^ and bit score of 200, and peptide data available from *D. melanogaster* database (FlyBase.org). Blast2GO was utilized to identify specific gene ontology (GO) terms that were enriched between treatments based on a Fisher’s Exact Test [50].

### Transmission electron microscopy

Cardia tissues from three non-inf, five inf+/- and six inf+/+ 40 day-old flies were dissected in 4% paraformaldehyde (PFA) and placed in 2.5% gluteraldehyde and 2% PFA in 0.1M sodium cacodylate buffer pH7.4 for 1 hour. Observed infected cardia were obtained from two different groups of flies independently infected with trypanosomes (n_1_=3 and n_2_=2 for inf+/-; n_1_=3 and n_2_=3 for inf+/+). Tissues were processed at the Yale Center for Cellular and Molecular Imaging (CCMI). Tissues were fixed in 1% osmium tetroxide, rinsed in 0.1M sodium cacodylate buffer and blocked and stained in 2% aqueous uranyl acetate for 1 hour. Subsequently, tissues were rinsed and dehydrated in a series in ethanol followed by embedment in resin infiltration Embed 812 (Electron Microscopy Sciences) and then stored overnight at 60°C. Hardened blocks were cut in sections at 60nm thickness using a Leica UltraCut UC7. The resulting sections were collected on formvar/carbon coated grids and contrast-stained in 2% uranyl acetate and lead citrate. Five grids including two sections prepared from each different insects were observed using a FEI Tecnai Biotwin transmission electron microscope at 80Kv. Images were taken using a Morada CCD camera piloted with the iTEM (Olympus) software. Contrasts of the pictures were adjusted using Photoshop CC 2018 (Adobe).

### Quantification of trypanosomes

At day 40 post parasite challenge, flies were dissected 72 hours after their last blood meal, and midgut and salivary glands (SG) were microscopically examined for infection status. Cardia were dissected, pooled by 5 in ice-cold TRIzol (Invitrogen) in function of their infection status (inf+/+ or inf+/-), and then flash-frozen in liquid nitrogen. RNA was extracted using the Direct-zol RNA MiniPrep (Zymo Research) following the manufacturer instructions, then subjected to DNase treatment using the Ambion TURBO DNA-free kit AM1907 (Thermo Fisher Scientific). 100ng of RNA was utilized to prepare cDNA using the iScript cDNA synthesis kit (Bio-Rad) following the manufacturer instructions. qPCR analysis was performed using SYBR Green supermix (Bio-Rad) and a Bio-Rad C1000 thermal cycler. Quantitative measurements were performed in duplicate for all samples. We used ATTCACGCTTTGGTTTGACC (forward) and GCATCCGCGTCATTCATAA (reverse) as primers to amplify trypanosome *gapdh*. We used CTGATTTCGTTGGTGATACT (forward) and CCAAATTCGTTGTCGTACCA (reverse) as primers to amplify tsetse *gapdh*. Relative density of parasite was inferred by normalizing trypanosome *gapdh* expression by tsetse *gapdh* expression. Statistical comparison of relative densities was performed on Prism 7 (GraphPad software) using a Student t-test.

Direct counting of parasites was operated by dissecting the cardia and the whole remaining midgut from flies prepared similarly than above. Individual tissues were homogenized in PSG buffer (8 replicates for each tissue). Homogenate was then fixed in an equal volume of 4% PFA for 30 min. The solution was then centrifuged 15 min at 110g, the supernatant was discarded and the pellets containing the trypanosomes from cardia and midguts were suspended in 100μl and 2,500μl PSG buffer, respectively. Trypanosomes from the total solution were counted using a hemocytometer. Statistical comparison of numbers was performed on Prism 7 (GraphPad software) using a Mann-Whitney rank test.

### Midgut-associated procyclic trypanosome re-infection

At day 40 post parasite challenge, flies were dissected 72 hours after their last blood meal, and midgut and salivary glands (SG) were microscopically examined for infection status. Around 40 inf+/+ and inf+/- were independently pooled together, and then roughly homogenized in 500μl of PSG buffer (PBS+2% glucose). Each homogenate was centrifuged 10 min at 30g to precipitate midgut debris, and then each supernatant containing parasites was transferred to a new tube to be centrifuged 15 min at 110g to precipitate the parasites. Supernatants were then discarded and each pellet containing midgut procyclic trypanosomes either from inf+/+ or inf+/- flies was suspended in 500μl PSG. Parasites were counted using a hemocytometer.

Newly emerged adult females were provided a blood diet including 10 μM Cysteine and supplemented with 5×10^6^ of procyclic trypanosomes from either inf+/+ or inf+/- flies prepared as described above. All flies were subsequently maintained on normal blood thereafter every 48 h. Four independent experiments were done for each type of trypanosomes. Midgut and salivary gland infections in each group were scored microscopically two weeks later. Precise sample sizes and count data are indicated in S1 Dataset.

Statistical analysis was carried out using the R software for macOS (version 3.3.2). A generalized linear model (GLM) was generated using binomial distribution with a logit transformation of the data. The binary infection status (inf+/+ or inf+/-) was analyzed as a function of the origin of the procyclic trypanosomes (inf+/+ or inf+/-) and the experiment it belongs to. The best statistical model was searched using a backward stepwise procedure from full additive model (*i.e*. parasite origin+experiment#) testing the main effect of each categorical explanatory factor. Using the retained model, we performed a Wald test on the individual regression parameters to test their statistical difference. Precise statistical results are indicated in S1 Dataset.

### Microscopic assessment of trypanosome developmental forms

Cardia from inf+/+ and inf+/- flies were dissected 40 dpa. Ten organs were pooled and gently homogenized in 100μL PBS and parasite numbers were evaluated using a hemocytometer. As cardia inf+/- contain less trypanosomes than cardia inf+/+, homogenates from cardia inf+/+ were diluted to the density of parasites present in cardia inf+/-. Equal numbers of parasites were then fixed in 2% Paraformaldehyde (PFA) PBS by adding an equal volume of 4% PFA PBS to the cardia inf+/+ and inf+/- homogenates. Parasites were then centrifuged for 10min at 500g and the resulting pellet was resuspended and washed in PBS. Samples were then centrifuged for 10min at 500g and the resulting pellet was resuspended in 200μL distilled water. 50μL of parasite-containing solution was deposited on poly-lysine coated slides and air dried. Slides were permeabilized for 10min in 0.1% Triton X-100 PBS, and then washed in PBS 5min and in distilled water 5min. Fluorescent DNA staining was then applied by covering the slides with a solution of DAPI in distilled water (1μg/mL) for 20 min in the dark. Slides were subsequently washed in distilled water two times for 5 min before being air dried in the dark. Microscopic observations were realized using a Zeiss AxioVision microscope (Zeiss). Detailed counts are indicated in S1 Dataset.

### VSG impact on PM-gene expression

Soluble VSG (sVSG) was prepared as described in [9]. Eight-day old adult flies received a blood meal containing purified sVSG (1μg/ml), or bovine serum albumin (BSA) (1μg/ml) as a control. To assess the effect of sVSG on gene expression at three days, cardia organs were microscopically dissected at 72h post treatment. To assess the effect of sVSG on gene expression at six days, remaining flies that were not dissected at three days were given a second normal blood meal, and the cardia organs were microscopically dissected at 72h post second feeding. Five biological replicates for each treatment and each time point were generated. Five dissected cardia were pooled for each replicate and their RNA was extracted. 100ng RNA was used to generate cDNA. Quantitative real-time PCR (qRT-PCR) was used to evaluate the expression of the PM-associated genes *proventriculin-1, −2* and *-3* as described in [9]. Normalization was performed to the internal control of *gadph* mRNA for each sample. Pairwise comparisons for each time point of the genes relative expression between sVSG and BSA treated flies was carried out with the Prism 7 software (GraphPad software) using a Student t-test.

Precise statistical results are indicated in S1 Dataset.

### RNAi-mediated knockdown of PM-associated gene expression

*Green fluorescent protein (gfp*) and *chitin synthase (cs*) gene specific dsRNAs were prepared as described in [7]. Newly emerged adult females were provided with a trypanosome supplemented blood diet that also included 10 μM Cysteine. All flies were subsequently maintained on normal blood thereafter every 48 h. After 6 days (at the 3rd blood meal), flies were divided into two treatment groups: first group received dsRNA-*cs* and the second group control dsRNA-*gfp*. The dsRNAs were administered to each group in 3 consecutive blood meals containing 3mg dsRNA/20μl blood (the approximate volume a tsetse fly imbibes each time it feeds). Four independent experiments using the same pool of dsRNA were generated for each treatment. Midgut and salivary gland infections in each group were scored microscopically three weeks later. Precise sample sizes and count data are indicated in S1 Dataset. Statistical analysis on the infection outcomes following the antioxidant feeding was carried out using the R software for macOS (version 3.3.2). A generalized linear model (GLM) was generated using binomial distribution with a logit transformation of the data. The binary infection status (inf+/+ or inf+/-) was analyzed as a function of the dsRNA treatment (dsRNA-*gfp* or dsRNA-cs) and the experiment it belongs to. The best statistical model was searched using a backward stepwise procedure from full additive model (*i.e*. dsRNA treatment+experiment#) testing the main effect of each categorical explanatory factors. Using the retained model, we performed a Wald test on the individual regression parameters to test their statistical difference. Precise statistical results are indicated in S1 Dataset.

Quantitative real-time PCR (qRT-PCR) was used to validate the effectiveness of our RNAi procedure as described in [7]. For each treatment of each experiment, we dissected the cardia of five randomly selected flies 72h after their third dsRNA-supplemented blood meal. The five dissected cardia were pooled together and their RNA was extracted. 100ng RNA was used to generate cDNA. RNA extractions from experiment #3 failed, but as the same dsRNA pools were used for all experiments and considering the consistency of the knockdown we observed, we decided to maintain experiment #3 in our counting results.

### *Serratia* infection assay to assess peritrophic matrix integrity

To assess the PM integrity, we applied a host survival assay following *per os* treatment of each group with *Serratia marcescens* as described in [7, 9]. We provided to three groups of 8 day-old flies (in their 4^th^ blood meal) either cardia extracts obtained from challenged flies that cleared the trypanosomes and are subsequently recovered from initial infection (rec-/-), or a cardia extract from inf+/- flies, or a cardia extract from inf+/+ flies. We included a fourth group of 8-day old flies that received an untreated blood meal.

Cardia extract was obtained by dissecting, in PBS, the cardia from 40 days-old infected as described above. Approximately fifty cardia from either rec-/-, inf+/- or inf+/+ flies were pooled together, and then gently homogenized. Parasites were counted from the homogenates of inf+/- and inf+/+ using a hemocytometer. The three cardia homogenates were then heated at 100°C for 10 minutes. inf+/- and inf+/+ extracts were provided to reach a concentration of 5×10^5^ parasites per ml of blood. As inf+/- cardia contain fewer parasites than inf+/+ cardia, the volume of the inf+/+ extract provided was adjusted by dilution in PSG buffer to be equal to inf+/- volume. Rec-/- extract was provided at an equal volume than infected extracts to ensure the presence of a similar quantity of extract molecules coming from the cardia in these groups. 48 hours after the flies received blood meal supplemented with the different extracts, all flies were provided a blood meal supplemented with 1,000 CFU/ml of *S. marcescens* strain Db11. Thereafter, flies were maintained on normal blood every other day, while their mortality was recorded every day for 30 days. Precise counting data are indicated in S1 Dataset.

Statistical analysis was carried out using the R software for macOS (version 3.3.2). We used an accelerated failure time model (Weibull distribution) where survival was analyzed as a function of the extract received (survreg() function of "survival" package). Pairwise tests were generated using Tukey contrasts on the survival model (glht() function of "multcomp" package). Precise statistical results are indicated in S1 Dataset.

### Antioxidant feeding

Newly emerged adult females were provided with a trypanosome-supplemented blood diet that also included 10 μM Cysteine. All flies were subsequently maintained on normal blood thereafter every 48 h. After 10 days (at the 5th blood meal), flies were divided into two treatment groups: first group received the anti-oxidant Cysteine (10μM) and the second group was fed normally as a control. Cysteine was administered each blood meal until dissection. Four independent experiments were done for each treatment. Midgut and salivary gland infections in each group were scored microscopically three weeks later. Precise sample sizes and count data are indicated in S1 Dataset.

Statistical analysis was carried out using the R software for macOS (version 3.3.2). A generalized linear model (GLM) was generated using binomial distribution with a logit transformation of the data. The binary infection status (inf+/+ or inf+/-) was analyzed as a function of the treatment (control or cysteine) and the experiment it belongs to. The best statistical model was searched using a backward stepwise procedure from full additive model (*i.e*. antioxidant treatment+experiment#) testing the main effect of each categorical explanatory factors. Using the retained model, we performed a Wald test on the individual regression parameters to test their statistical difference. Precise statistical results are indicated in S1 Dataset.

### Quantification of reactive oxygen species (ROS) in cardia tissues

ROS were quantified using the Amplex Red Hydrogen Peroxide/Peroxidase Assay Kit (ThermoFisher Scientific), following the manufacturer recommendations. 40 days post parasite challenge, flies were dissected 72 hours after their last blood meal, and midgut and salivary glands (SG) were microscopically examined for infection status. For each infection phenotype (*i.e*. inf+/+ or inf+/-), 3 replicates containing each 10 cardia tissues pooled and homogenized in 80μl of ice-cold Amplex Red Kit 1X Reaction Buffer were generated. Three replicates of age-matched non-infected cardia tissues were conceived in the same manner. 50μl of assay reaction mix was added to 50μl of the supernatant of each samples, and then incubated 60 minutes at RT. Fluorescence units were determined using a BioTek Synergy HT plate reader (530nm excitation; 590nm emission). Peroxide concentrations were determined using the BioTek Gen5 software calculation inferred from a standard curve (precise results are indicated in S1 Dataset). Statistical analysis was performed on Prism 7 (GraphPad software) using a one-way ANOVA where ROS concentration was analyzed as a function of the infection status. Pairwise comparisons were carried out using a TukeyHSD posthoc test.

## Acknowledgements

We thank Dr. Ying Yang for technical assistance with the tsetse fly colony, Drs. Nikolay Kolev and Chris Tschudi for their advice with data interpretation, Morven Graham for her assistance with electron microscopy data interpretation, and Drs. Maria Onyango, Mehmet Karakus and Florent Masson for their critical comments on the manuscript.

## Supporting Information

### S1 Fig. Overview of cardia transcriptomes

(**A**) Number of RNA-seq reads in each of three biological replicates from Non-inf, Inf +/- and Inf +/+ cardia. (**B**) Proportion of total trimmed reads that map to *Glossina morsitans morsitans* or *Trypanosoma brucei brucei 927*. (**C**) Percent relative abundance of mapped *Glossina morsitans morsitans* transcripts.

### S2 Fig. Parasite quantity in midgut and cardia

(**A**) Number of parasites in the cardia of Inf+/- (red) and Inf+/+ (blue) flies. (**B**) Number of parasites in the midgut of Inf+/- (red) and Inf+/+ (blue) flies. The black bar represents the mean of the replicates for each treatment. Midgut and cardia were dissected from eight 40 days-old females. Parasites were counted using a hemocytometer. Statistical analyses were carried out using the non-parametrical Mann-Whitney rank test.

### S3 Fig. Ultrastructure of the cardia

(**A**) Transversal section of a non-infected cardia. Two pictures of the same cardia were merged to produce a larger picture. B, C, D, E and F are magnified micrographs of cardia tissues. (**B**) Midgut tissue delimiting the outer part of the cardia. (**C**) Foregut tissue invagination within cardia, corresponding to the stomodeal valve in other insects. (**D**) Myofibrils assembled to form the sphincter surrounding the foregut opening in the cardia. (**E**) Lipid-containing cells, immediately adjacent to foregut tissue, covered by a thin layer of muscle. (**F**) Foregut tube coming out of the cardia. LU: Lumen; MG: Midgut; PM sc: PM secreting cells; FG: Foregut; MU: muscle; LC: Lipid-containing cells; mv: microvilli; ES: Ecotperitrophic space.

### S4 Fig. Expression of *chitin synthase* after RNAi treatment

Expression of *chitin synthase* relative to constitutively expressed *β-tubulin* after treatment with dsRNA-gfp (control; white circles) and dsRNA-*chitin synthase* (dsCS; grey circles). *chitin synthase* expression is significantly decreased after RNAi knockdown (Student t-test, p = 0.011).

### S5 Fig. Ultrastructure of trypanosomes in cardia inf+/+

Right panels present accumulation of parasites next to the PM secreting cells in the cardia. Left panels present trypanosomes next to mature PM. Micrographs in this image represent four of six biological replicates. Right and left panels are paired to correspond to a same individual. Blue arrowheads: newly secreted PM; red arrowhead: PM electron-dense layer; green arrowhead: PM electron-lucent layer; MG: midgut; PM sc: PM secreting cells; mv: microvilli; TR: trypanosomes; LU: lumen; FG: foregut; MU: muscle.

### S6 Fig. Ultrastructure of cardia from inf+/+ tsetse, showing trypanosomes inbetween layers of the PM

(**A**) Ultrastructure of the cardia in the region where the PM is secreted. Trypanosomes are observed either trapped inbetween the electron-dense layer of the PM (red arrowhead) and newly synthesized PM secretions (blue arrowheads), or free in the lumen and close to the foregut (yellow arrowhead). (**B**) is a magnified micrograph of the black frame in (A). (**C**) Ultrastructure of the region below the annular cleft where the PM is secreted. Trypanosomes are observed in the ES, in the lumen and trapped in the PM as cyst-like bodies (purple arrowheads). (**D**) Magnified micrograph of the white frame in (A). A cyst-like body (purple arrowhead) is entrapped inbetween the electron-dense (red arrowhead) and electron-lucent layers of the PM (green arrowhead). The ultrastructure presented in micrographs (A) and (C-D) originated from 2 different inf+/+ cardia. Micrographs in this image represent 2 two of six of biological replicates from cardia inf+/+. MG: midgut; PM sc: PM secreting cells; mv: microvilli; TR: trypanosomes; LU: lumen; FG: foregut; MU: muscle; ES: ectoperitrophic space.

### S7 Fig. Ultrastructure of cardia and PM from non-inf and inf+/+ tsetse

(**A**) PM ultrastructure in cardia of non-inf tsetse. The electron-dense (red arrowhead) and electron-lucent (green arrowhead) layers of a mature PM are intact in these flies (**B-D**) Close-up images of disrupted electron-dense layer of the PMs. Cyst-like bodies (purple arrowheads) are observed inbetween the two layers of the PM. Micrographs in this image represent one and three of three and six of biological replicates from cardia non-inf and inf+/+, respectively. PM: peritrophic matrix; MG: midgut; mv: microvilli; TR: trypanosomes; LU: lumen; ES: ectoperitrophic space.

### S8 Fig. Differentially expressed (DE) transcripts in parasitized cardia

(**A**) A total of 2,093 transcripts were DE in inf+/- and inf+/+ cardia relative to uninfected (non-inf) controls. In inf+/- cardia (red), 429 and 327 transcripts were up and downregulated, respectively. In inf+/+ cardia (blue), 278 and 413 transcripts were up and downregulated, respectively. Of the DE transcripts shared by both infection phenotypes (646), 576 are similarly regulated, while 70 show DE in inf+/- versus inf+/+ phenotypes. (**B-C**) Transcripts are plotted as a function of their fold-changes (Log2 scale) obtained by comparison between control non-inf transcriptome and either inf+/+ (y-axis) or inf+/- (x-axis) transcriptome. The size of the circle indicates the expression value (CPM) in the control non-inf cardia. The genes presented have been annotated with their genome ID number and their best BLASTx annotation. Panel B shows similarly regulated transcripts in both cardia infection phenotypes. Panel C shows differentially regulated transcripts between the two cardia infection phenotypes. In panel C, DE transcripts expressed exclusively in inf+/+ compared to non-inf are shown in blue, DE transcripts expressed exclusively in inf+/- compared to non-inf are shown in red, and DE transcripts expressed in both inf+/+ and inf+/- compared to non-inf are shown in green.

### S9 Fig. Ultrastructure of muscles and mitochondria in cardia from inf+/- tsetse

(**A**) Transversal section of muscle tissues (MU) composing the sphincter. (**B-D**) Longitudinal section of muscles (MU) layering the midgut tissues. Micrographs in this image represent three of five of biological replicates from cardia inf+/-. Black arrowheads: healthy mitochondria; red arrowheads: vacuolation of mitochondria; yellow arrow: sarcoplasmic dilatation; green arrowheads: swelling mitochondria; MG: midgut.

---

**S1 Dataset. Detailed results and statistics for infection experiments and oxidative stress quantification.**

**S2 Dataset. Detailed results and analyses fo each transcriptome.**

**S3 Dataset. GO terms analysis results.**

## References

1. Simarro PP, Cecchi G, Franco JR, Paone M, Diarra A, Priotto G, et al. Monitoring the progress towards the elimination of gambiense human African trypanosomiasis. PLOS Negl Trop Dis. 2015;9(6):e0003785. Epub 2015/06/10. doi: 10.1371/journal.pntd.0003785. PubMed PMID: 26056823; PubMed Central PMCID: PMCPMC4461311.

2. Simarro PP, Cecchi G, Franco JR, Paone M, Diarra A, Ruiz-Postigo JA, et al. Estimating and mapping the population at risk of sleeping sickness. PLOS Negl Trop Dis. 2012;6(10):e1859. Epub 2012/11/13. doi: 10.1371/journal.pntd.0001859. PubMed PMID: 23145192; PubMed Central PMCID: PMCPMC3493382.

3. Courtin F, Camara M, Rayaisse JB, Kagbadouno M, Dama E, Camara O, et al. Reducing human-tsetse contact significantly enhances the efficacy of sleeping sickness active screening campaigns: a promising result in the context of elimination. PLOS Negl Trop Dis. 2015;9(8):e0003727. Epub 2015/08/13. doi: 10.1371/journal.pntd.0003727. PubMed PMID: 26267667; PubMed Central PMCID: PMCPMC4534387.

4. Davis S, Aksoy S, Galvani A.A global sensitivity analysis for African sleeping sickness. Parasitology. 2011;138(4):516–26. Epub 2010/11/17. doi: 10.1017/S0031182010001496. PubMed PMID: 21078220; PubMed Central PMCID: PMCPMC3282146.

5. Hegedus D, Erlandson M, Gillott C, Toprak U. New insights into peritrophic matrix synthesis, architecture, and function. Annu Rev Entomol. 2009;54:285–302. Epub 2008/12/11. doi: 10.1146/annurev.ento.54.110807.090559. PubMed PMID: 19067633.

6. Rose C, Belmonte R, Armstrong SD, Molyneux G, Haines LR, Lehane MJ, et al. An investigation into the protein composition of the teneral *Glossina morsitans morsitans* peritrophic matrix. PLOS Negl Trop Dis. 2014;8(4):e2691. Epub 2014/04/26. doi: 10.1371/journal.pntd.0002691. PubMed PMID: 24763256; PubMed Central PMCID: PMCPMC3998921.

7. Weiss BL, Savage AF, Griffith BC, Wu Y, Aksoy S. The peritrophic matrix mediates differential infection outcomes in the tsetse fly gut following challenge with commensal, pathogenic, and parasitic microbes. J Immunol. 2014;193(2):773–82. Epub 2014/06/11. doi: 10.4049/jimmunol.1400163. PubMed PMID: 24913976; PubMed Central PMCID: PMCPMC4107339.

8. Rodgers FH, Gendrin M, Wyer CAS, Christophides GK. Microbiota-induced peritrophic matrix regulates midgut homeostasis and prevents systemic infection of malaria vector mosquitoes. PLOS Pathog. 2017;13(5):e1006391. Epub 2017/05/26. doi:10.1371/journal.ppat.1006391. PubMed PMID: 28545061; PubMed Central PMCID: PMCPMC5448818.

9. Aksoy E, Vigneron A, Bing X, Zhao X, O’Neill M, Wu YN, et al. Mammalian African trypanosome VSG coat enhances tsetse’s vector competence. Proc Natl Acad Sci U S A. 2016;113(25):6961–6. Epub 2016/05/18. doi: 10.1073/pnas.1600304113. PubMed PMID: 27185908; PubMed Central PMCID: PMCPMC4922192.

10. Coutinho-Abreu IV, Sharma NK, Robles-Murguia M, Ramalho-Ortigao M. Characterization of *Phlebotomus papatasi* peritrophins, and the role of PpPer1 in *Leishmania* major survival in its natural vector. PLOS Negl Trop Dis. 2013;7(3):e2132. Epub 2013/03/22. doi: 10.1371/journal.pntd.0002132. PubMed PMID: 23516661; PubMed Central PMCID: PMCPMC3597473.

11. Narasimhan S, Rajeevan N, Liu L, Zhao YO, Heisig J, Pan J, et al. Gut microbiota of the tick vector *Ixodes scapularis* modulate colonization of the Lyme disease spirochete. Cell Host Microbe. 2014;15(1):58–71. Epub 2014/01/21. doi: 10.1016/j.chom.2013.12.001. PubMed PMID: 24439898; PubMed Central PMCID: PMCPMC3905459.

12. Turner CM, Barry JD, Vickerman K. Loss of variable antigen during transformation of *Trypanosoma brucei rhodesiense* from bloodstream to procyclic forms in the tsetse fly. Parasitol Res. 1988;74(6):507–11. Epub 1988/01/01. PubMed PMID: 3194363.

13. Hao Z, Kasumba I, Lehane MJ, Gibson WC, Kwon J, Aksoy S. Tsetse immune responses and trypanosome transmission: implications for the development of tsetse-based strategies to reduce trypanosomiasis. Proc Natl Acad Sci U S A. 2001;98(22):12648–53. Epub 2001/10/11. doi: 10.1073/pnas.221363798. PubMed PMID: 11592981; PubMed Central PMCID: PMCPMC60108.

14. Hu Y, Aksoy S. An antimicrobial peptide with trypanocidal activity characterized from *Glossina morsitans morsitans*. Insect Biochem Mol Biol. 2005;35(2):105–15. Epub 2005/02/01. doi: 10.1016/j.ibmb.2004.10.007. PubMed PMID: 15681221.

15. MacLeod ET, Maudlin I, Darby AC, Welburn SC. Antioxidants promote establishment of trypanosome infections in tsetse. Parasitology. 2007;134(Pt 6):827–31. Epub 2007/02/20. doi: 10.1017/S0031182007002247. PubMed PMID: 17306056.

16. Wang J, Aksoy S. PGRP-LB is a maternally transmitted immune milk protein that influences symbiosis and parasitism in tsetse’s offspring. Proc Natl Acad Sci USA. 2012;109(26):10552–7. Epub 2012/06/13. doi: 10.1073/pnas.lll6431109. PubMed PMID: 22689989; PubMed Central PMCID: PMCPMC3387098.

17. Haines LR, Lehane SM, Pearson TW, Lehane MJ. Tsetse EP protein protects the fly midgut from trypanosome establishment. PLOS Pathog. 2010;6(3):e1000793. Epub 2010/03/12. doi: 10.1371/journal.ppat.1000793. PubMed PMID: 20221444; PubMed Central PMCID: PMCPMC2832768.

18. Sharma R, Peacock L, Gluenz E, Gull K, Gibson W, Carrington M. Asymmetric cell division as a route to reduction in cell length and change in cell morphology in trypanosomes. Protist. 2008;159(1): 137–51. Epub 2007/10/13. doi: 10.1016/j.protis.2007.07.004. PubMed PMID: 17931969.

19. Van Den Abbeele J, Claes Y, van Bockstaele D, Le Ray D, Coosemans M. *Trypanosoma brucei* spp. development in the tsetse fly: characterization of the post-mesocyclic stages in the foregut and proboscis. Parasitology. 1999;118 ( Pt 5):469–78. Epub 1999/06/11. PubMed PMID: 10363280.

20. Peacock L, Ferris V, Bailey M, Gibson W. Dynamics of infection and competition between two strains of *Trypanosoma brucei brucei* in the tsetse fly observed using fluorescent markers. Kinetoplastid Biol Dis. 2007;6:4. Epub 2007/06/08. doi: 10.1186/1475-9292-6-4. PubMed PMID: 17553128; PubMed Central PMCID: PMCPMC1899512.

21. Haines LR. Examining the tsetse teneral phenomenon and permissiveness to trypanosome infection. Front Cell Infect Microbiol. 2013;3:84. Epub 2013/12/07. doi: 10.3389/fcimb.2013.00084. PubMed PMID: 24312903; PubMed Central PMCID: PMCPMC3833344.

22. Welburn SC, Maudlin I. The nature of the teneral state in *Glossina* and its role in the acquisition of trypanosome infection in tsetse. Ann Trop Med Parasitol. 1992;86(5):529–36. Epub 1992/10/01. PubMed PMID: 1288435.

23. Aksoy S, Gibson WC, Lehane MJ. Interactions between tsetse and trypanosomes with implications for the control of trypanosomiasis. Adv Parasitol. 2003;53:1–83. Epub 2003/11/01. PubMed PMID: 14587696.

24. Dyer NA, Rose C, Ejeh NO, Acosta-Serrano A. Flying tryps: survival and maturation of trypanosomes in tsetse flies. Trends Parasitol. 2013;29(4):188–96. Epub 2013/03/20. doi: 10.1016/j.pt.2013.02.003. PubMed PMID: 23507033.

25. Buxton P. The natural history of tsetse flies. London: H. K. Lewis and Co. Ltd.; 1955.

26. Lehane MJ. Peritrophic matrix structure and function. Annu Rev Entomol. 1997;42:525–50. Epub 1997/01/01. doi: 10.1146/annurev.ento.42.1.525. PubMed PMID: 15012322.

27. Wigglesworth VB. Digestion in the tsetse-fly: a study of structure and function. Parasitology. 1929;21(3):288–321.

28. Moloo SK, Kutuza SB. Feeding and crop emptying in *Glossina brevipalpis* Newstead. Acta Trop. 1970;27(4):356–77. Epub 1970/01/01. PubMed PMID: 4396362.

29. Moloo SK, Steiger RF, Hecker H. Ultrastructure of the peritrophic membrane formation in *Glossina* Wiedemann. Acta Trop. 1970;27(4):378–83. Epub 1970/01/01. PubMed PMID: 4101510.

30. Freeman JC. The penetration of the peritrophic membrane of the tsetse flies by trypanosomes. Acta Trop. 1973;30(4):347–55. Epub 1973/04/01. PubMed PMID: 4149681.

31. Steiger RF. On the ultrastructure of *Trypanosoma (Trypanozoon) brucei* in the course of its life cycle and some related aspects. Acta Trop. 1973;30(1):64–168. Epub 1973/01/01. PubMed PMID: 4144959.

32. Hao Z, Aksoy S. Proventriculus-specific cDNAs characterized from the tsetse, *Glossina morsitans morsitans*. Insect Biochem Mol Biol. 2002;32(12):1663–71. Epub 2002/11/14. PubMed PMID: 12429118.

33. Gibson W, Bailey M. The development of *Trypanosoma brucei* within the tsetse fly midgut observed using green fluorescent trypanosomes. Kinetoplastid Biol Dis. 2003;2(1):1. Epub 2003/05/29. PubMed PMID: 12769824; PubMed Central PMCID: PMCPMC156611.

34. Schuster S, Krüger T, Subota I, Thusek S, Rotureau B, Beilhack A, et al. Developmental adaptations of trypanosome motility to the tsetse fly host environments unravel a multifaceted *in vivo* microswimmer system. eLife. 2017;6:e27656. doi: 10.7554/eLife.27656.

35. Kuraishi T, Binggeli O, Opota O, Buchon N, Lemaitre B. Genetic evidence for a protective role of the peritrophic matrix against intestinal bacterial infection in *Drosophila melanogaster*. Proc Natl Acad Sci U S A. 2011;108(38):15966–71. Epub 2011/09/08. doi: 10.1073/pnas. 1105994108. PubMed PMID: 21896728; PubMed Central PMCID: PMCPMC3179054.

36. Lehane MJ, Allingham PG, Weglicki P. Composition of the peritrophic matrix of the tsetse fly, *Glossina morsitans morsitans*. Cell Tissue Res. 1996;283(3):375–84. Epub 1996/03/01. PubMed PMID: 8593667.

37. Oberholzer M, Lopez MA, McLelland BT, Hill KL. Social motility in African trypanosomes. PLOS Pathog. 2010;6(l):e1000739. Epub 2010/02/04. doi: 10.1371/journal.ppat.1000739. PubMed PMID: 20126443; PubMed Central PMCID: PMCPMC2813273.

38. Imhof S, Knusel S, Gunasekera K, Vu XL, Roditi I. Social motility of African trypanosomes is a property of a distinct life-cycle stage that occurs early in tsetse fly transmission. PLOS Pathog. 2014;10(10):e1004493. Epub 2014/10/31. doi: 10.1371/journal.ppat.1004493. PubMed PMID: 25357194; PubMed Central PMCID: PMCPMC4214818.

39. Attardo GM, Strickler-Dinglasan P, Perkin SA, Caler E, Bonaldo MF, Soares MB, et al. Analysis of fat body transcriptome from the adult tsetse fly, *Glossina morsitans morsitans*. Insect Mol Biol. 2006;15(4):411–24. Epub 2006/08/16. doi: 10.1111/j.1365-2583.2006.00649.x. PubMed PMID: 16907828.

40. Murphy MP. How mitochondria produce reactive oxygen species. Biochem J. 2009;417(1):1–13. Epub 2008/12/09. doi: 10.1042/BJ20081386. PubMed PMID: 19061483; PubMed Central PMCID: PMCPMC2605959.

41. Kowaltowski AJ, Vercesi AE. Mitochondrial damage induced by conditions of oxidative stress. Free Radic Biol Med. 1999;26(3-4):463–71. Epub 1999/01/23. PubMed PMID: 9895239.

42. Wang Y, Nartiss Y, Steipe B, McQuibban GA, Kim PK. ROS-induced mitochondrial depolarization initiates PARK2/PARKIN-dependent mitochondrial degradation by autophagy. Autophagy. 2012;8(10):1462–76. Epub 2012/08/15. doi: 10.4161/auto.21211. PubMed PMID: 22889933.

43. Bravard A, Bonnard C, Durand A, Chauvin MA, Favier R, Vidal H, et al. Inhibition of xanthine oxidase reduces hyperglycemia-induced oxidative stress and improves mitochondrial alterations in skeletal muscle of diabetic mice. Am J Physiol Endocrinol Metab. 2011;300(3):E581–91. Epub 2011/01/13. doi: 10.1152/ajpendo.00455.2010. PubMed PMID: 21224483.

44. Ridgley EL, Xiong ZH, Ruben L. Reactive oxygen species activate a Ca2+-dependent cell death pathway in the unicellular organism *Trypanosoma brucei brucei*. Biochem J. 1999;340 ( Pt l):33–40. Epub 1999/05/07. PubMed PMID: 10229656; PubMed Central PMCID: PMCPMC1220219.

45. Ha EM, Oh CT, Ryu JH, Bae YS, Kang SW, Jang IH, et al. An antioxidant system required for host protection against gut infection in Drosophila. Dev Cell. 2005;8(l):125–32. Epub 2004/12/29. doi: 10.1016/j.devcel.2004.11.007. PubMed PMID: 15621536.

46. Molina-Cruz A, DeJong RJ, Charles B, Gupta L, Kumar S, Jaramillo-Gutierrez G, et al. Reactive oxygen species modulate *Anopheles gambiae* immunity against bacteria and *Plasmodium*. J Biol Chem. 2008;283(6):3217–23. Epub 2007/12/11. doi: 10.1074/jbc.M705873200. PubMed PMID: 18065421.

47. Van Den Abbeele J, Caljon G, De Ridder K, De Baetselier P, Coosemans M. *Trypanosoma brucei* modifies the tsetse salivary composition, altering the fly feeding behavior that favors parasite transmission. PLOS Pathog. 2010;6(6):e1000926. Epub 2010/06/10. doi: 10.1371/journal.ppat.1000926. PubMed PMID: 20532213; PubMed Central PMCID: PMCPMC2880569.

48. Peacock L, Ferris V, Bailey M, Gibson W. The influence of sex and fly species on the development of trypanosomes in tsetse flies. PLoS Negl Trop Dis. 2012;6(2):e1515. Epub 2012/02/22. doi: 10.1371/journal.pntd.0001515. PubMed PMID: 22348165; PubMed Central PMCID: PMCPMC3279344.

49. Robinson MD, McCarthy DJ, Smyth GK. edgeR: a Bioconductor package for differential expression analysis of digital gene expression data. Bioinformatics. 2010;26(l):139–40. Epub 2009/11/17. doi: 10.1093/bioinformatics/btp616. PubMed PMID: 19910308; PubMed Central PMCID: PMCPMC2796818.

50. Conesa A, Gotz S, Garcia-Gomez JM, Terol J, Talon M, Robles M. Blast2GO: a universal tool for annotation, visualization and analysis in functional genomics research. Bioinformatics. 2005;21(18):3674–6. Epub 2005/08/06. doi: 10.1093/bioinformatics/bti610. PubMed PMID: 16081474.

